# Disrupting microglial TGF-β signaling triggers region-specific pathology in the spinal cord

**DOI:** 10.1101/2023.04.24.538074

**Authors:** Keying Zhu, Jin-Hong Min, Vijay Joshua, Yun Liu, Melanie Pieber, Valerie Suerth, Heela Sarlus, Robert Harris, Harald Lund

**Affiliations:** Applied Immunology and Immunotherapy, Department of Clinical Neuroscience, Karolinska Institutet, Center for Molecular Medicine, Karolinska University Hospital, Stockholm, Sweden; Division of Rheumatology, Department of Medicine Solna, Karolinska Institutet, Karolinska University Hospital, Stockholm, Sweden; MOE Key Laboratory of Metabolism and Molecular Medicine, Department of Biochemistry and Molecular Biology, School of Basic Medical Sciences and Zhongshan Hospital, Fudan University, Shanghai, China; State Key Laboratory of Medical Neurobiology and MOE Frontiers Center for Brain Sciences, Institutes of Brain Science, Fudan University, Shanghai, China; Department of Physiology and Pharmacology, Karolinska Institutet, Center for Molecular Medicine, Karolinska University Hospital, Stockholm Sweden

## Abstract

Transforming growth factor-β (TGF-β) signaling is critical for microglial maturation during development and the maintenance of microglial homeostasis in adulthood. It remains unclear whether regional susceptibilities to the loss of TGF-β signaling in microglia also exist, and the contributing factors have yet to be identified. We find that deletion of *Tgfbr2* on microglia leads to microglial activation and demyelination in mouse spinal cords, primarily in the dorsal column (DC). *Tgfbr2*-deficient microglia exhibit distinct transcriptomic changes, and those sorted from the DC display a more proinflammatory profile compared to those from the ventral column (VC) and grey matter (GM). Single nucleus RNA sequencing (snRNA-seq) of the spinal cord uncovers a microglial subtype that emerges exclusively following *Tgfbr2* deletion (termed TGFβ signaling-suppressed microglia, TSM), exhibiting high expression of *Mmp12, Gpnmb, Lgals3, Mgll, and Alcam,* predominantly located in the DC. Phenotypically, disruption of microglial TGF-β signaling results in behavioral deficits that are more severe in female and older mice, whereas young male mice are less affected. Mechanistically, we reveal a significantly higher level of TGF-β1/TGFBR2 in the spinal cords of normal older mice compared to the young mice, with the DC region richer in genes of the TGF-β signaling pathway than the VC and GM regions. This indicates that older mice and the DC region require more TGFβ1 to maintain tissue homeostasis and, reciprocally, are more responsive and sensitive to the disruption of TGF-β signaling in microglia. Herein, we report a demyelinating disease with region-specificity and its susceptibility to the loss of microglial TGF-β signaling with gender and age differences. Our findings contribute valuable information to our understanding of the importance of microglia in regulating myelin health, especially during the aging process.

## Introduction

One of the most important signaling pathways regulating microglial development, maturation, and homeostasis is the transforming growth factor-β (TGF-β) signaling pathway. Although not critical for microglial survival, TGF-β signaling is essential for adopting microglial identity during early development and for the maintenance of a functional signature of microglia^1,2^. TGF-β signaling is crucial for the developmental distinction between microglia and border associated macrophages (BAMs) in the central nervous system (CNS); loss of TGF-β signaling in microglia affects their embryonic expansion and alters their phenotype, leading to downregulation of microglial signature genes (such as *P2ry12*) and upregulation of BAM genes (such as *Mrc1*), as well as a proinflammatory profile^1^. However, the importance of TGF-β signaling in adult microglia and how it choreographs homeostasis in the adult CNS is not entirely understood. Whether TGF-β signaling is critical for microglia-specific signature in adult microglia is still controversial, as a previous study using the *Cx3cr1^CreERT2^Tgfbr2^fl/fl^* mice to delete Tgfbr2 in adult microglia demonstrate no change in microglia-specific gene signature (such as *Pr2y12, Sall1, Olfml3, and Tmem119*)^3,4^ or behavioral impairments. This argues against our previous findings and other studies^5–7^.

We previously used the genetic *Cx3cr1^CreER/+^ R26^DTA/+^* mouse tool to deplete microglia followed by chimeric reconstitution of monocytes lacking *Tgfbr2* (*LysM ^Cre/+^Tgfbr2 ^fl/fl^→ Cx3cr1^CreER/+^ R26^DTA/+^*) and found that TGF-β signaling is required for monocytes to enter the CNS and functionally integrate into the empty microglial niche^6^. Engrafted monocyte-derived macrophages in the microglia-depleted CNS, with disrupted TGF-β signaling, failed to adopt a microglia-like signature as the wild-type macrophages do, leading to a fatal disorder with demyelinating pathologies primarily in the dorsal and dorsolateral columns of spinal cord. However, it remained to be confirmed whether microglial *Tgfbr2* deletion leads to a similar demyelinating pattern since seminal studies have proven that monocyte-derived macrophages and CNS resident microglia have distinct origins and may be functionally different^8–13^. Consistent with our observation^6^, dorsal column-specific demyelinating pathology has also been reported in another study using mice lacking the milieu molecule LRRC33^14^. This molecule is mainly expressed in microglia and acts as an essential ‘anchoring’ receptor for the activation of latent TGF-β^14^. Nevertheless, the molecular understanding responsible for the demyelinating pathology specific to the dorsal region remains elusive. We therefore hypothesized that the microenvironment in the subregions of the spinal cord may influence microglial sensitivity to the loss of TGF-β signaling.

In this study, we comprehensively examined the role of TGF-β signaling in adult microglia, and disclosed the cellular and molecular mechanisms, along with the age and gender influences, that drive the spinal cord region-specific demyelinating disease induced by microglia with impaired TGF-β signaling. Our findings deepen the understanding of how age-related changes in the spinal cord microenvironment render microglial dependence on TGF-β signaling, and further highlight the important role of microglia in regulating myelin health, especially in aging conditions.

## Results

### Loss of *Tgfbr2* induces aberrant microglial responses and demyelinating pathology in the dorsal column of the spinal cord

To understand the function of TGF-β signaling in adult microglia, we employed the *Cx3cr1^CreER^Tgfbr2^fl/fl^* mice, in which tamoxifen administration induces *Tgfbr2* deletion in *Cx3cr1*-expressing cells. Our previous study revealed that when CNS microglia were replaced by monocyte-derived macrophages lacking *Tgfbr2*, demyelinating pathologies and myelin-ingesting foamy macrophages appeared in the spinal cords. Here we also found that following microglial *Tgfbr2* deletion, enhanced immunoreactivity of Iba1 and MHC-II was observed mostly in the dorsal column (DC), whereas the ventral column (VC) remained less affected (Fig. 1 a-c); dorsolateral columns also showed increased Iba1 and MHC-II expression at later stages. Fluoromyelin staining in the spinal cord revealed myelin loss in the DC but not in the VC, starting from around 30 days after microglial *Tgfbr2* deletion, and microglia in the DC were laden with ingested myelin fragments (Fig. 1 d-f). Microglia within the demyelinated DC area exhibited a typical foamy morphology (Fig. 1 d, f), and those at the edge of the DC were more ramified yet were interacting with myelin fragments found in close proximity to microglia processes (Fig. 1 f). To analyze microglial activation in the different spinal cord subregions, we analyzed microglia isolated from micro-dissected DC, VC, and grey matter (GM) of the spinal cord using flow cytometry. We found that microglial F4/80 expression increased significantly after 18 days of *Tgfbr2* deletion and remained at a high level after 28 days, although its expression at D28 decreased compared to that at the D18 time point (Fig. 1 g, h). MHC-II and CD45 expression increased over time following the deletion of *Tgfbr2* and were also significantly higher than that in the wild-type (WT) microglia (Fig 1. g, h). The percentage of MHC-II^+^ and CD45^+^ microglia was also higher in the DC compared to other spinal cord regions (Fig 1. i, j), whereas the percentage of F4/80^+^ microglia across the three subregions was similar (Fig. 1 k), possibly because the increase in F4/80 expression occurred much earlier than the other surface markers. These data confirm that disruption of microglial TGF-β signaling in adult mice triggers aberrant microglial response, leading to demyelinating pathology in the dorsal column of the spinal cord.

**Fig 1.**
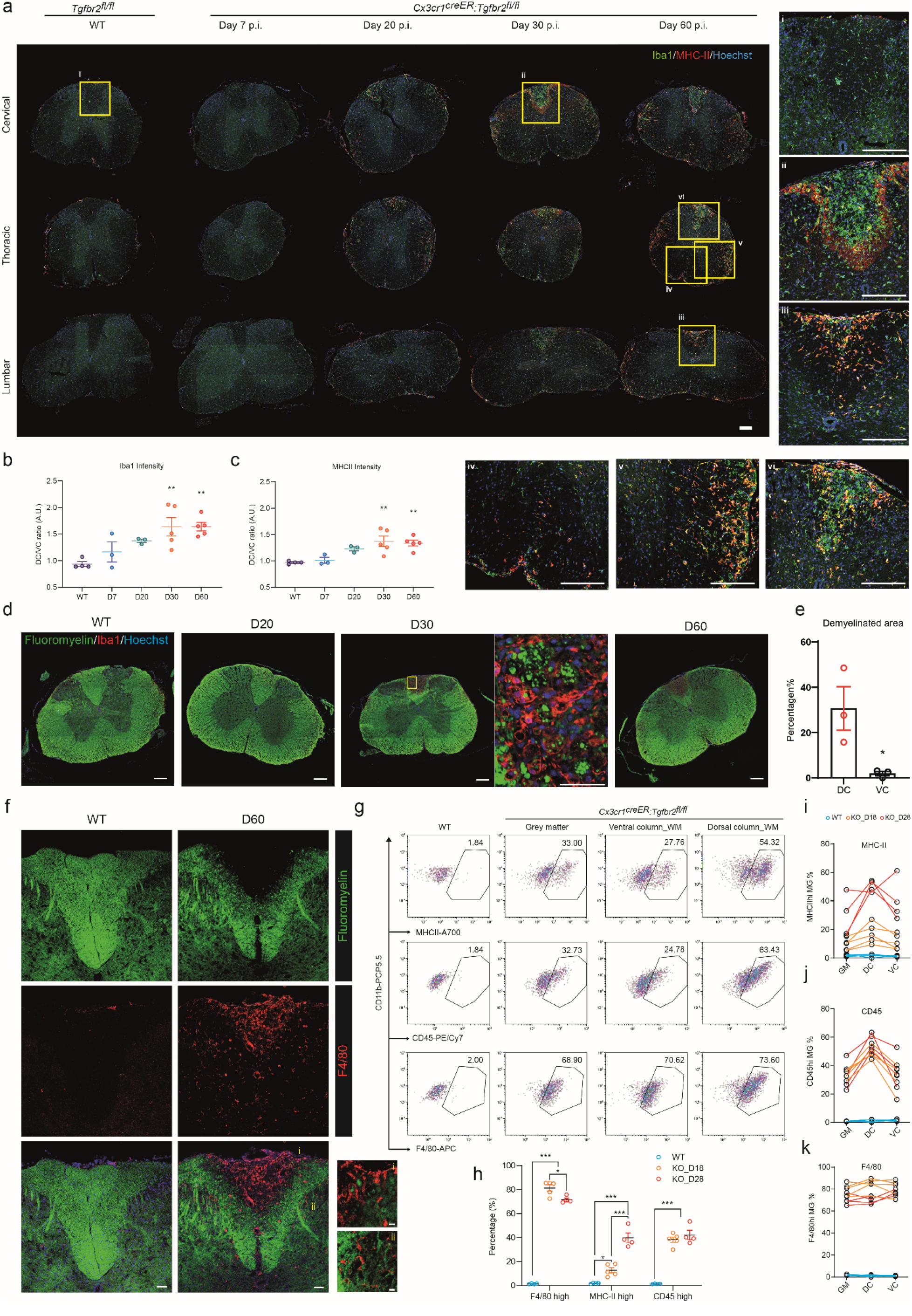
Aberrant microglial states and demyelination in the spinal cord in mice deficient of microglial *Tgfbr2*. **a**, Representative immunohistostaining of adult female (4-7 months) spinal cord sections from cervical, thoracic and lumbar segments of the *Cx3cr1^creER^:Tgfbr2^fl/fl^* mice 7, 20, 30, and 60 days post tamoxifen injection and the age-paired wild-type (WT) *Tgfbr2^fl/fl^* control mice (Iba1 in green, MHC-II in red, and nuclei staining in blue). Insets i-vi are the yellow boxes indicated in the main image. Scale bar, 200 μm. **b,c,** The fluorescent intensity ratio of Iba1 **(b)** and MHC-II **(c)** between the dorsal column white matter and the ventral column white matter of the thoracic segments. n = 3-5 mice per group. A.U., arbitrary units. **d**, Representative immunohistostaining of spinal cord sections from cervical segments of the *Cx3cr1^creER^:Tgfbr2^fl/fl^* mice 20, 30, and 60 days post tamoxifen injection and the WT mice (Fluoromyelin in green, Iba1 in red, and nuclei staining in blue). Scale bar, 200 μm. The inset of the yellow box in the D30 section is attached to its right showing myelin ingestion by Iba1+ foamy cells in the dorsal column white matter (DC). Scale bar, 50 μm. **e**, The percentage of demyelinated area in the respect DC or ventral column white matter (VC). n = 3 mice per group. **f**, Representative immunohistostaining of Fluoromyelin (green) and F4/80 (red) in the DC showing demyelination and microglial activation. Scale bar, 200 μm. Insets of the indicated regions (i) and (ii) in the D60 section are attached with (i) showing F4/80+ foamy cells with ingested myelin inside the demyelinating area, and (ii) showing ramified F4/80+ cells interacting with myelin at the edge of DC. Scale bar for the insets, 10 μm. **g**, Representative flow cytometry dot plots of CD11b^+^CD45^+^Ly6C^-^ cells (microglia) microdissected from the DC, VC and grey matter (GM) regions from a 4-month-old female *Cx3cr1^creER^:Tgfbr2^fl/fl^* mouse 28 days post tamoxifen injection and an age-paired female wild-type mouse (microglia from the dorsal column white matter were used as a control reference in the WT panel). The numbers above the indicated gates represent the percentage (%) of the gated cells in the total CD11b^+^CD45^+^Ly6C^-^ cells. **h**, Quantification relating to (g) showing the frequency of F4/80 high, MHC-II high, and CD45 high cells in the total CD11b^+^CD45^+^Ly6C^-^ cells among the WT mice, and *Cx3cr1^creER^:Tgfbr2^fl/fl^* mice 18 and 28 days after tamoxifen injection. Cells from the DC, VC, and GM regions of the same mouse were pooled for analyses. n = 4 mice per group. **i, j, k**, Line charts relating to (g) showing the cross-regional change in the frequency of MHC-II high (i), CD45 high (j), and F4/80 high (k) cells. Data are shown as mean ± SEM. \**p*⍰<⍰0.05; \*\**p*⍰<⍰0.01; \*\*\**p*⍰<⍰0.001. Unpaired Student’s t-test was used in (e); One-way ANOVA with Dunnett’s Multiple Comparison Test was used in (b) and (c) to compare the WT group with other groups; one-way ANOVA with Tukey’s Multiple Comparison Test was used in (h).

### Spinal cord transcriptomic analyses reveal global molecular changes in the absence of microglial *Tgfbr2*

To better unravel the global molecular changes in the spinal cords of mice with microglial *Tgfbr2* deletion, we employed the NanoString Glial Profiling Panel to analyze a set of curated (∽780) genes related to glial cell function, neuroinflammation, and different cell states. The spinal cord tissues from wild-type (WT) mice and *Cx3cr1^CreER^Tgfbr2^fl/fl^* mice after 10 (KO-D10), 20 (KO-D20), and 30 (KO-D30) days of tamoxifen administration were analyzed (Fig. 2 a). We selected these time points to analyze the molecular changes occurring in the spinal cord during the acute phase of microglial *Tgfbr2* deletion (D10), before the onset of demyelinating pathology (D20), and during the onset of demyelination (D30). Unbiased principal component (PC) analysis revealed that the spinal cords of the microglial *Tgfbr2*- deleted mice were already distinct from those of the WT mice after 10 days of microglial *Tgfbr2* deletion, and this segregation remained evident for the D20 and D30 samples as well (Fig. 2 b). The volcano plot presented the differentially expressed genes (DEGs) between the KO and WT spinal cords, revealing the downregulation of microglia-specific genes *P2ry12* and *Cx3cr1* and upregulation of genes involved in phagocytosis (*Msr1*), antigen presentation (such as *H2-d1 and Cd74*), and microglia neurodegenerative phenotype (MGnD) or disease-associated microglia (DAM) (such as *Trem2, Apoe, Lpl, Itgax, and Ctsb*) (Fig. 2 c, j; Supplementary Fig. 1 b, c). Interestingly, *Tgfb1* and *Nrros* (encoding LRRC33), involved in the TGF-β signaling pathway, were also found upregulated in the microglial *Tgfbr2*-deleted spinal cords, indicating a potential compensatory mechanism to facilitate a favorable milieu for TGF-β signaling transduction when proper TGF-β signaling in microglia was disrupted (Fig. 2 c; Supplementary Fig 1. d-i).

**Fig 2.**
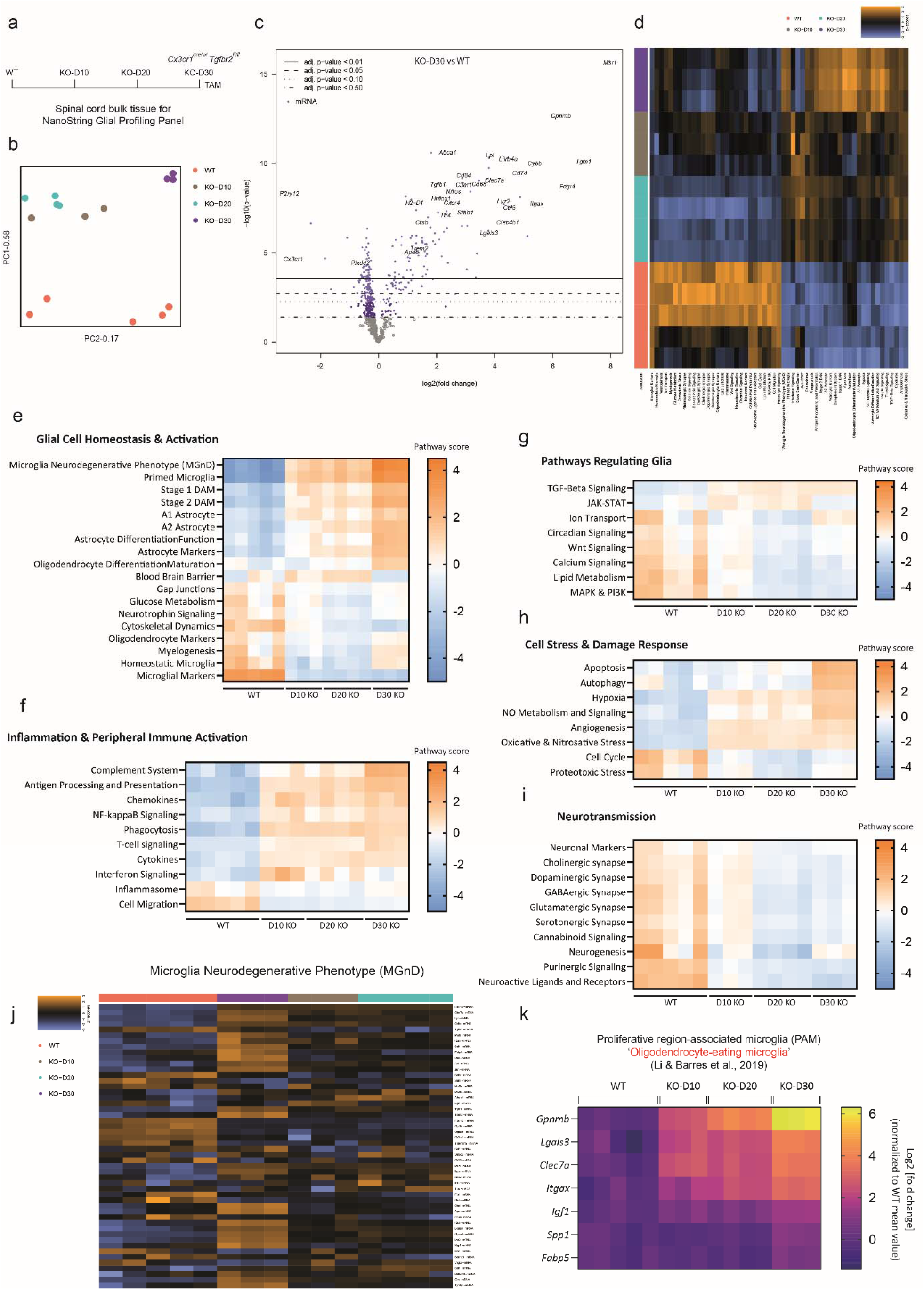
Profiling of molecular signatures in the spinal cord after microglial deletion of *Tgfbr2*. **a**, Schematic illustration of the experiment: spinal cords from 14-17-month-old female wild-type (WT) and *Cx3cr1^creER^:Tgfbr2^fl/fl^* mice mice (KO-D10, KO-D20, KO-D30 denotes 10, 20, and 30 days post tamoxifen injection) were homogenized and total RNA were extracted for a NanoString Glial Profiling Panel analysis covering 780 genes across different biological pathways involved in glial cell biology. **b**, Unbiased principal component (PC) analysis of based on the normalized mRNA counts of all samples; *n* = 3-5 samples per group. **c**, Volcano plots of mRNA count data displaying each gene’s -log10(p-value) and log2 fold change (*Tgfbr2^-/-^* KO-D30/WT). Horizontal lines indicate various False Discovery Rate (FDR) thresholds or p-value thresholds if there is no adjustment to the p-values. Genes are colored if the resulting p-value is below the given FDR or p-value threshold. **d**, Heatmap of pathway scores presenting a high-level overview of how the pathway scores change across samples. High scores are in orange and low scores are in blue. Scores are displayed on the same scale via a Z-transformation. **e-i**, NanoString pathway scores were presented using heatmap to show how pathways involved in glial cell homeostasis and activation (e), inflammation and peripheral immune activation (f), glial cell regulation (g), cell stress and damage response (h), and neurotransmission (i) vary across groups. Each square indicates the score of an indicated pathway of one sample; high scores are in orange, and lows scores are in blue. **j**, Heatmap with unsupervised clustering of normalized mRNA counts of genes scaled to give all genes equal variance in the pathway *Microglia Neurodegenerative Phenotype (MGnD)*; orange indicates high expression, and blue indicates low expression. **k**, Heatmap of the normalized mRNA expression (Log2 transformed) of genes identified by Li & Barres et al. as key signature genes of proliferative region-associated microglia (PAM) were plotted across samples.

The curated genes included in the NanoString panel were annotated with different molecular functions and pathways (Supplementary Fig. 1 a), and the heatmap depicting the clustering of the spinal cord samples across the pathways (based on the pathway scores given by NanoString) showed that the WT and microglial *Tgfbr2*-deleted spinal cords were enriched in very distinct molecular pathways (Fig. 2 d). Among all the molecular pathways, the ones that are highly relevant to microglial function, such as *Microglial Markers, Microglia Neurodegenerative Phenotype (MGnD), and Primed Microglia,* showed striking changes chronologically, and pathways involved in neurotransmission were globally suppressed following microglial *Tgfbr2* deletion, indicating a potential inhibition of normal neuronal activities in the spinal cords (Fig. 2 e-j). Oligodendrocyte markers (included in the panel: *Mbp, Mog, Plp1, Sox10, Ugt8, and Ermn*), together with the *Myelogenesis* pathway which indicates the formation of myelin sheath, were downregulated at D10 and D20 KO groups as compared to the WT group. However, we noted a slight enrichment of the *Myelogenesis* pathway and upregulated expression of genes involved in oligodendrocyte differentiation and maturation at D30, possibly suggesting an active differentiating process of oligodendrocyte progenitors to compensate for the loss of myelin at this timepoint. Activated and neuroinflammatory microglia are known to trigger neurotoxic reactive astrocytes, which can be prevented by TGF-β^15^. Accordingly, we also found that disrupting microglial TGF-β signaling induces astrocyte reactivity (Fig. 2 e). In addition, inflammatory signatures, such as *NF-kappaB Signaling, Cytokines, Complement System, Antigen Processing and Presentation*, and signatures related to cell stress and damage, such as Apoptosis and NO Metabolism and Signaling, were induced in the spinal cords after deletion of *Tgfbr2* in microglia, which were indicative of inflammatory responses and dysregulated cellular function in the spinal cords (Fig. 2 f, h).

A previous study identified a microglial subtype called proliferative region-associated microglia (PAMs) in the developing white matter of postnatal mice (P7), which phagocytose newly formed oligodendrocytes^16^. PAMs have similar gene signatures to DAMs/MGnDs that are observed in neurodegenerative conditions in adult mice, but the top signature genes for PAMs are *Gpnmb* and *Spp1*, and PAM induction is independent of the TREM2-APOE pathway. Since we also found that microglia are ingesting myelin in the absence of *Tgfbr2*, we wondered if the signature genes of PAMs were also altered. We plotted the mRNA expression of the signature genes of PAMs among our spinal cord samples and found that the expression of PAM signature genes, such as *Gpnmb, Lgals3, and Clec7a*, increased over time after microglial *Tgfbr2* deletion (Fig. 2 k). This increase corresponded well with the occurrence of the demyelinating pathology that was prominent after 30 days of microglial *Tgfbr2* deletion (Fig. 1 d), suggesting an association of the PAM genes with the ongoing demyelinating pathology in the spinal cord. Collectively, these data show that microglial deficiency of *Tgfbr2* triggers extensive molecular changes in the spinal cords.

### *Tgfbr2*-deleted microglia adopt altered gene signature

We next wondered what transcriptomic processes were happening in these *Tgfbr2*-deficient adult microglia. Since we also noticed differential microglial responses in the different subregions in spinal cords, we therefore sorted microglia from the DC, VC, and GM regions from WT and microglial *Tgfbr2*- deleted mice after 18 (D18) or 28 (D28) days of tamoxifen injection and performed bulk RNA-sequencing (RNA-seq) on the sorted cells (Fig. 3 a). We first pooled the microglia from the three regions to compare the transcriptomic changes between the WT and *Tgfbr2*-deleted microglia. We identified 668 DEGs when comparing the D18 *Tgfbr2*-deleted microglia with the WT microglia, and 585 DEGs between the D28 *Tgfbr2*-deleted and WT microglia (Fig.3 b). 416 genes remained differentially expressed at both time points when compared to the WT microglia, and we named them core DEGs. We plotted the top 50 hub genes of the core DEGs and their regulatory network using *Cytohubba*^17^. Among the induced hub genes, we noted prominent proinflammatory gene signature (such as *Il1b* and *Vcam1*), antigen presentation signature (such as *H2-Ab1*), and chemokine activities (such as *Ccl7, Ccl8, Cxcl1, Cxcl2, Cxcl10, Cxcl12, Cxcr3, and Cxcr4*), indicating that these microglia were highly activated and being recruited (Fig.3 c, d). Apart from the inflammatory signature, we also found that genes encoding C-type lectins and MMPs were upregulated after *Tgfbr2* deletion, suggesting that these microglia may actively interact with other cells by remodeling the ECM (Fig. 3 d). MS4A genes, primarily expressed by myeloid cells, were also upregulated (Fig. 3 d). Much of the function of MS4A genes are unknown, but they are implicated in regulating macrophage function^18^. *Ms4a4a* expression in macrophages is induced by IL-4 stimulation, and it facilitates the activation of Dectin-1 receptor and Syk-dependent signaling pathway, implying a role in pattern-recognition and endocytosis^19–21^. Interestingly, we also noted induced expression of genes (*Abca1, Apoe*, and *Plin2*) involved in cholesterol metabolism (Fig. 3 d), a metabolic process that plays a decisive role for microglia to process lipid-rich myelin debris and recycle cholesterol for oligodendrocytes to make new myelin^22–24^. Of note, unlike the previous report^3^, we found that among the core DEGs, microglia-specific gene signatures were globally suppressed after microglial *Tgfbr2* deletion (Fig. 3 d), including *Cx3cr1, Tmem119, Sall1, Olfml3, Hexb, Siglech,* and *Sparc,* proving that disruption of TGF-β signaling in adult microglia affects their homeostatic states. Kyoto Encyclopedia of Genes and Genomes (KEGG) pathway analysis and Biological Process Gene Ontology (GO_P) analysis revealed major transcriptomic changes in inflammatory responses, innate immune responses, antigen processing/presentation, and chemotaxis (Fig. 3 e, f). Concomitant with the identified DEGs, key pathways that were relevant to the observed microglial-driven demyelinating pathology following *Tgfbr2* deletion, such as *Cytokine-cytokine receptor interaction, Cell adhesion molecules (CAMs), TNF signaling pathway, Phagosome, Chemokine signaling pathway, and Cholesterol metabolism,* were highly enriched after microglial deletion of *Tgfbr2* (Fig. 3 e, f).

**Fig 3.**
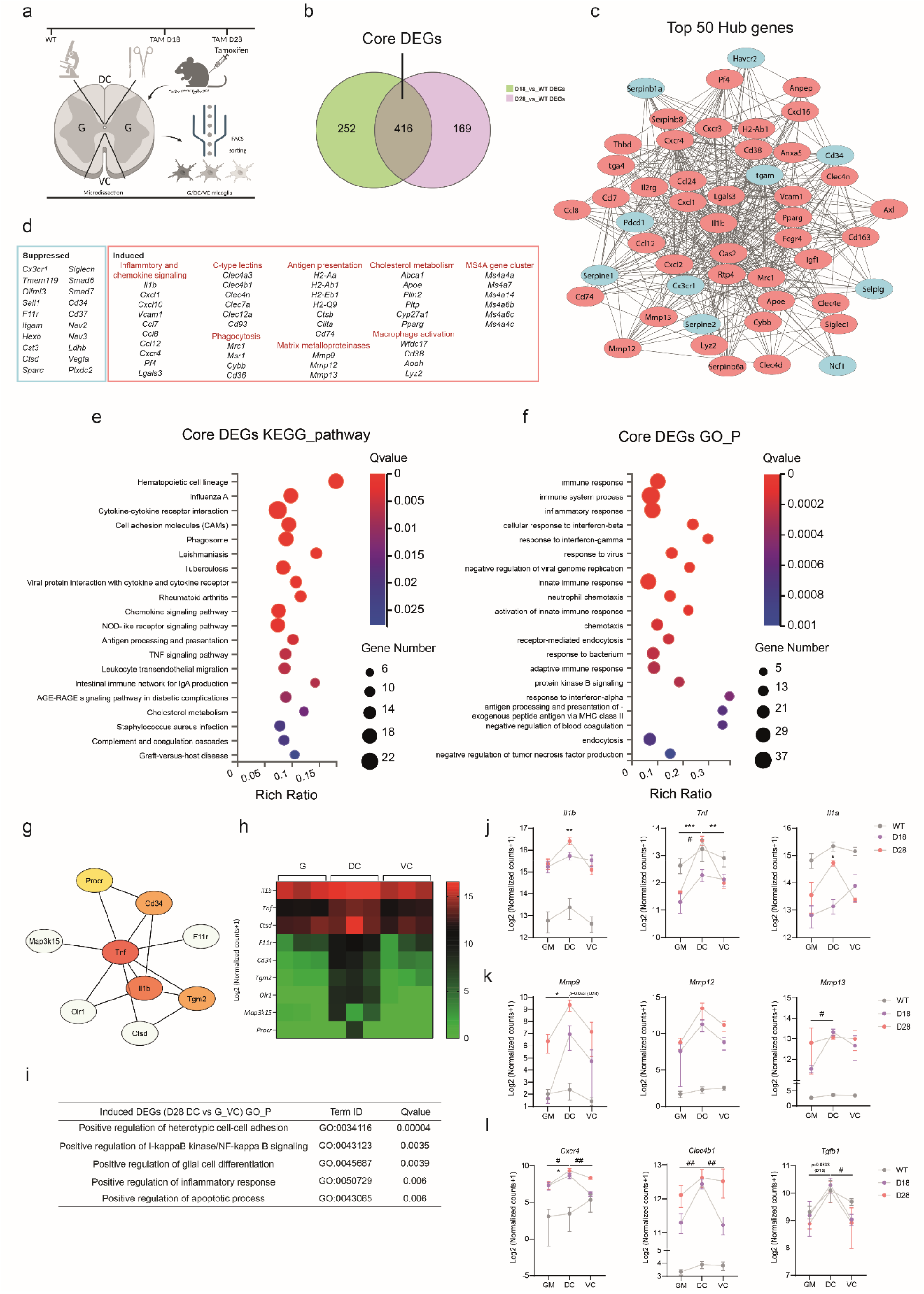
Transcriptomic analysis of microglia sorted from different spinal cord regions. **a**, Schematic representation of the experimental design: spinal cords from wildtype mice (WT) and tamoxifen injected *Cx3cr1^creER^:Tgfbr2^fl/fl^* mice (18 and 28 days post injection, referred to as D18 and D28, respectively) were microdissected under a dissecting microscope to separate the dorsal column white matter (DC), ventral column white matter (VC), and the grey matter (GM). The dissected tissues were processed individually to obtain single-cell suspensions followed by flow cytometry sorting for microglia. 100 sorted microglia from each region of each mouse were then processed for bulk-RNA sequencing. **b**, Venn diagram showing the number of the differentially expressed genes (DEGs) between the D18 and WT groups (D18_vs_WT: 668 DEGs), and between the D28 and WT groups (D28_vs_WT: 585 DEGs), and the overlapping 416 genes, which are named the core DEGs. DEGs were identified after DESeq2 normalization (Log2 fold change ≥ 2, qvalue < 0.01). **c**, Top 50 hub genes from the 416 core DEGs and their interacting network were revealed with Cytohubba using the MCC algorithm. Genes in red bubbles are upregulated genes after microglial *Tgfbr2* depletion (compared to the WT group), and blue bubbles are downregulated genes. **d**, Selected suppressed genes and induced gene clusters following microglial *Tgfbr2* deletion compared to WT microglia. **e, f** Kyoto Encyclopedia of Genes and Genomes (KEGG) pathway enrichment analysis and Gene Ontology Biological Process (GO_P) enrichment analysis of the 416 core DEGs (performed using Dr.Tom analysis tool based on the phyper function in R software; qvalue was obtained by correction of pvalue, and qvalue ≤ 0.05 is regarded as a significant enrichment). The bubble size reflects the number of DEGs. The colors of the bubbles illustrate the q values for each term (low in red, and high in blue). ‘Rich ratio’ indicates the ratio of Term Candidate Gene Number versus the total Term Gene Number. **g**, The 30 DEGs identified by comparing microglia from DC with microglia from VC and G from D28 mice (DESeq2 normalized; Log2 fold change ≥1, qvalue < 0.05) were analyzed using Cytohubba with the MCC algorithm and the hub genes and their interactions were presented. The colored bubbles are the top 5 hub genes (descending order from red, orange to yellow). **h**, Heatmap showing the expression of the DEGs (Log2 transformed) in (g) among the DC, VC and GM regions. **i**, The enriched GO_P terms of the induced DEGs comparing microglia from DC with microglia from VC and GM from D28 mice. **j, k, l** Cross-regional representation of selected genes, including pro-apoptotic genes (j), metalloproteinases genes (k), and other genes of interest (l) from the 416 core DEGs in (b). Data are shown as mean ± SEM. # denotes comparison among D18 samples (compared to D18 DC), and * denotes comparison among D28 samples (compared to D28 DC). # or *, *p* < 0.05; ## or **, *p* < 0.01; ### or ***, *p* < 0.001; One-way ANOVA with Dunnett’s Multiple Comparison Test.

To further understand the regional heterogeneity of microglia in the spinal cord, we next compared the transcriptomic change of microglia sorted from the DC, VC, and GM regions. Under a homeostatic condition in the WT mice, no transcriptomic differences were identified among the three regions (data not shown), which was consistent with previous findings showing that homeostatic microglia in adult CNS show limited transcriptomic heterogeneity^16^. When comparing the D28 DC microglia with the D28 VC and GM microglia, we identified 30 DEGs. Although the number of the DEGs was small, they were highly involved in inflammatory process, pro-apoptotic activities, cell-cell adhesion, and glial cell function, which may underlie the microglia-myelin interaction happening in the DC (Fig. 3 g-i). Temporal and spatial comparison of cytokine genes that were detrimental to oligodendrocyte survival (*Il1b, Tnf, Il1a*)^25–29^, as well as matrix metalloproteinase genes (*Mmp9, Mmp12, Mmp13*), which were involved in degrading ECM^30^ and myelin proteins^31–33^ and processing latent TGF-β complex^34,35^, revealed a higher induction in the DC microglia (Fig.3 j, k). We also noticed a higher expression of *Cxcr4* and *Clec4b1* in the DC microglia, and interestingly, *Tgfb1* expression remained higher in the DC microglia irrespective of the existence of *Tgfbr2* or not (Fig. 3 l). Taken together, these data reveal transcriptomic changes of adult spinal cord microglia after *Tgfbr2* deletion with subregional information.

### snRNA-seq reveals a subcluster of microglia appearing in the dorsal column after *Tgfbr2* deletion

Enzymatic digestion and dissociation used for microglial isolation and subsequent RNA-seq can lead to significant artifacts and loss of microglial transcriptomic information due to the ability of microglia to sense and quickly adapt to changes in the microenvironment ^36–38^. To minimize these artificial effects and gain a better understanding of the involvement of different cell types in the spinal cord demyelinating pathology after *Tgfbr2* deletion in microglia, we utilized single nucleus RNA sequencing (snRNA-seq) on freshly dissected spinal cord tissues. This approach involved snap-freezing and mechanical dissociation of the tissue to obtain nuclei for sequencing analyses, allowing for a more accurate representation of microglial gene expression.

Through uniform manifold approximation and projection (UMAP) clustering followed by cell cluster annotation, we identified major cell types in the spinal cords, including subtypes of neurons, motor neurons, subtypes of oligodendrocytes and myelin, astrocytes, microglia, and pericytes (Fig.4 a; Supplementary Fig. 2 a). We did not find a major contribution of meningeal cells as the other dataset reported^39^, mainly because we flushed the spinal cord tissues out from the spine and the meninges were removed in this way. No peripheral immune cell populations were identified, indicating deleting microglial *Tgfrb2* did not trigger peripheral immune cell infiltration to the spinal cord, which was consistent with our previous findings^6^. We noted dynamic changes primarily in the proportion of glial cells following 20 days (D20) and 30 days (D30) of microglial *Tgfbr2* deletion, but no significant alterations in the neuron subtypes (Fig. 4 b; Supplementary Fig. 2 b). Specifically, the microglia population (MG) enlarged over time, whereas the myelin/mature oligodendrocyte population (Myelin_OL) disappeared to a large degree. Interestingly, after microglial *Tgfbr2* deletion, we observed a cluster of OPCs (OPC-2) that proliferated, while a cluster of oligodendrocytes (OL-1) enlarged at D20 but returned to normal or lower levels at D30. Another oligodendrocyte cluster (OL-2) initially decreased at D20 but recovered its population at D30. We speculate that the OL-2 cluster may represent a later stage of oligodendrocyte differentiation, and this OPC-2_→_OL-1_→_OL-2 oligodendrocyte lineage differentiation may serve as a compensatory process to address the loss of mature oligodendrocytes (Myelin_OL) or OL-2 due to microglial *Tgfbr2* deletion. Despite the impact on the population of Myelin_OL and OL-2 at D20, OL-2 was able to recover by D30, whereas the population of Myelin_OL remained unchanged. This suggests that the loss of *Tgfrb2* in microglia did not affect the early stages of OPC/oligodendrocyte differentiation, but rather hindered the ability of highly differentiated oligodendrocytes to myelinate.

**Fig 4.**
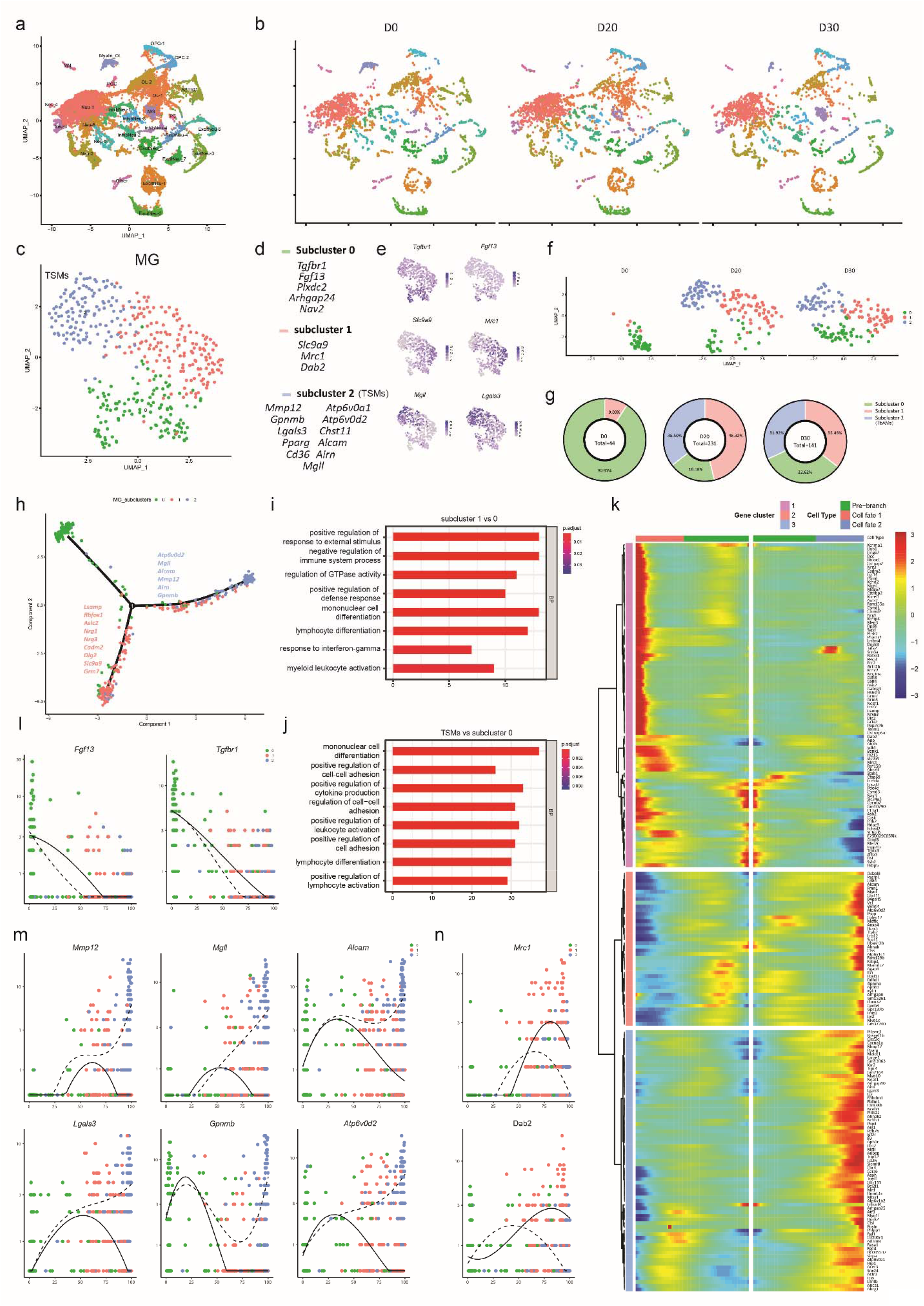
Single nucleus RNA-sequencing (snRNA-seq) reveals induced microglial subclusters after depletion of *Tgfbr2* in microglia. **a**, UMAP of 26, 053 pooled spinal cord cells from 12-month-old female wild-type (D0) and Cx3cr1^CreER^:Tgfbr2^fl/fl^ mice after 20 days (D20) and 30 days (D30) of tamoxifen injection, with annotation of each cluster. **b**, UMAP of the cells from the D0, D20, and D30 groups (2500 cells for each group) showing the change of each cluster overtime. c, UMAP of 416 cells in the microglia (MG) cluster from (a). MG subcluster 0 in green, MG subcluster 1 in red, and MG subcluster 2 in blue. **d**, Signature genes of the MG subclusters identified in (c). **e**, Representative signature gene expression across the 3 MG subclusters. **f**, UMAP of MG cells from the D0, D20, and D30 groups showing the change of each MG subcluster overtime. There are 44, 231, and 141 MG in D0, D20, and D30 groups, respectively; 44 MG are plotted for D0 group, and 141 MG are plotted for D20 and D30 groups. **g**, The percentage of each MG subcluster for each time point with the total number of MG cells indicated in the middle. **h**, Pseudotime trajectory of MG cells differentiating from MG subcluster 0 to either MG subcluster 1 or MG subcluster 2 (or TSM) with determinant genes for each cell fate at the branching point. **i, j,** Gene Ontology analysis of DEGs identified between MG subcluster 1 and MG subcluster 0 (i), or DEGs identified between MG subcluster 2 (or TSM) and MG subcluster 0 (j). **k**, Heatmap showing genes that are branch-dependent expressed in the branch point of the pseudotime trajectory of MG cells. Columns are points in pseudotime, rows are genes, and the beginning of pseudotime is in the middle of the heatmap. Cell fate 1 represents the lineage of MG subcluster 1, and cell fate 2 represents the lineage of MG subcluster 2 (or TSM). **l**, The pseudotime expression of selected genes downregulated from MG subcluster 0 to MG subcluster 1 and 2. **m**, The pseudotime expression of selected determinant genes for MG subcluster 2 (or TSM). **n**, The pseudotime expression of selected determinant genes for MG subcluster 1.

The microglia population was further clustered into three subclusters (Fig. 4 c-e). The signature genes of subcluster 0 included *Tgfbr1, Fgf13*, and *Plxdc2*, and the majority of the microglia at D0 (naive state) belonged to this subcluster. The proportion of subcluster 0 decreased after *Tgfbr2* deletion, indicating that this subcluster may represent homeostatic microglia, despite the absence of typical microglia-specific signatures such as *P2ry12* and *Tmem119*, which may be due to the potential technical limitation of snRNA-seq (Fig.4 f, g). The signature genes of subcluster 1 were *Slc9a9, Mrc1*, and *Dab2*, and very few cells of subcluster 1 existed at D0, but its population expanded after *Tgfbr2* deletion (Fig.4 d-g). It has been reported that microglial *Mrc1* expression increases after the inhibition of TGF-β signaling, and *Dab2* encodes a specific adaptor protein that prevents the degradation of *Tgfbr2* and facilitates the transduction of TGF-β1/SMAD signaling^40–42^. In addition, *Mrc1* and *Dab2* were highly expressed by border associated macrophages, and therefore subcluster 1 may also include CNS macrophages^43,44^. What we found the most interesting was subcluster 2, which did not exist at the naïve state (D0) and was induced only after disrupting microglial TGF-β signaling (Fig.4 f, g), indicating that this population was highly relevant and specific to the loss of the TGF-β signaling. We therefore denoted this subcluster 2 MG as TGF-β signaling-suppressed microglia (TSM). TSMs express signature genes such as *Mmp12, Gpnmb, Lgals3, Alcam, Mgll*, and *Pparg* (Fig.4, d, e). We further conducted a pseudo-time trajectory analysis on the MG population and verified that the cells differentiated from subcluster 0 to either subcluster 1 or TSMs (Fig.4 h). The determinant genes for differentiating into subcluster 1 at the branching point were *Lsamp, Rbfox1, Asic2, Nrg1, Nrg3, Cadm2, Dlg2, Slc9a9,* and *Grm7*, and the determinant genes for TSMs were *Atp6v0d2, Mgll, Alcam, Mmp12, Airn, and Gpnmb*. GO biological process analyses revealed that compared to subcluster 0, subcluster 1 was enriched in pathways involved in immune cell differentiation and activation, GTPase activity and response to external stimulus and interferon-gamma, whereas TSMs were enriched in pathways regulating cytokine production, cell-cell adhesion, and immune cell differentiation and activation (Fig. 4 i, j). We plotted the genes that were expressed in a branch-dependent manner in the branch point of the pseudotime trajectory of MG cells, and this revealed gene clusters corresponding to different cell states (Fig. 4 k). Specifically, we noted that genes such as *Fgf13* and *Tgfbr1* were downregulated over the pseudotime differentiation from the pre-branch state (subcluster 0) to the other cell states (Fig. 4 l). Genes including *Mmp12, Mgll, Alcam, Lgals3, Gpnmb*, and *Atp6v0d2* were upregulated in the cell fate 2 (TSMs) when differentiating from subcluster 0 (Fig. 4 m), and *Mrc1* and *Dab2* were mainly induced in the cell fate 1 (subcluster 1) during the differentiation (Fig. 4 n).

Of note, the appearance of TSMs (20 days after microglial *Tgfbr2* deletion) preceded the demyelinating pathology in the DC, implying that they may play an active role in the progression of DC demyelination rather than simply serving as scavengers for myelin debris. On the other hand, the subcluster 1 MG, due to enriched expression of *Mrc1* and *Dab2*, genes known to be neuroprotective in neurodegenerative or neuroinflammatory conditions^41,45–47^, may be induced to restrain the damage. Coincidentally, the two signature genes of TSMs, *Gpnmb* and *Lgals3*, were also the key signature genes of the abovementioned PAMs that phagocytose newly formed oligodendrocytes in the postnatal white matter^16^. Additionally, our NanoString analyses (Fig. 2 c, k) also revealed a chronological enrichment of *Gpnmb* and *Lgals3* in the spinal cord after microglial *Tgfbr2* deletion. We therefore were interested in the spatial distribution of these TSM signature genes in the spinal cord. Immunostaining revealed a significant increase in the expression of GPNMB and Galectin-3 (or MAC-2; encoded by *Lgals3*) in the spinal cord of microglial *Tgfbr2*-deleted mice (Fig. 5 a-g). Furthermore, the expression of these markers was found to be significantly higher in the DC compared to the VC (Fig. 5 a-g). We also observed higher expression of Galectin-3 and GPNMB in the lateral columns, where mild demyelination was also evident. Co-staining with the microglia/macrophage marker F4/80 and Iba1 confirmed the colocalization of GPNMB and Galectin-3 with microglia displaying foamy morphology in the DC (Fig.5 d, h), indicating that the induced TSMs were mainly located in the DC region, highlighting their engagement in the DC demyelinating pathology. Collectively, these data reveal that glial cells in the spinal cords, but not the neurons, were highly affected following microglial *Tgfbr2* deletion and that a TGF-β signaling-suppressed microglia subtype (TSM) were highly induced in the DC region and may be involved in the disruption of myelin.

**Fig 5.**
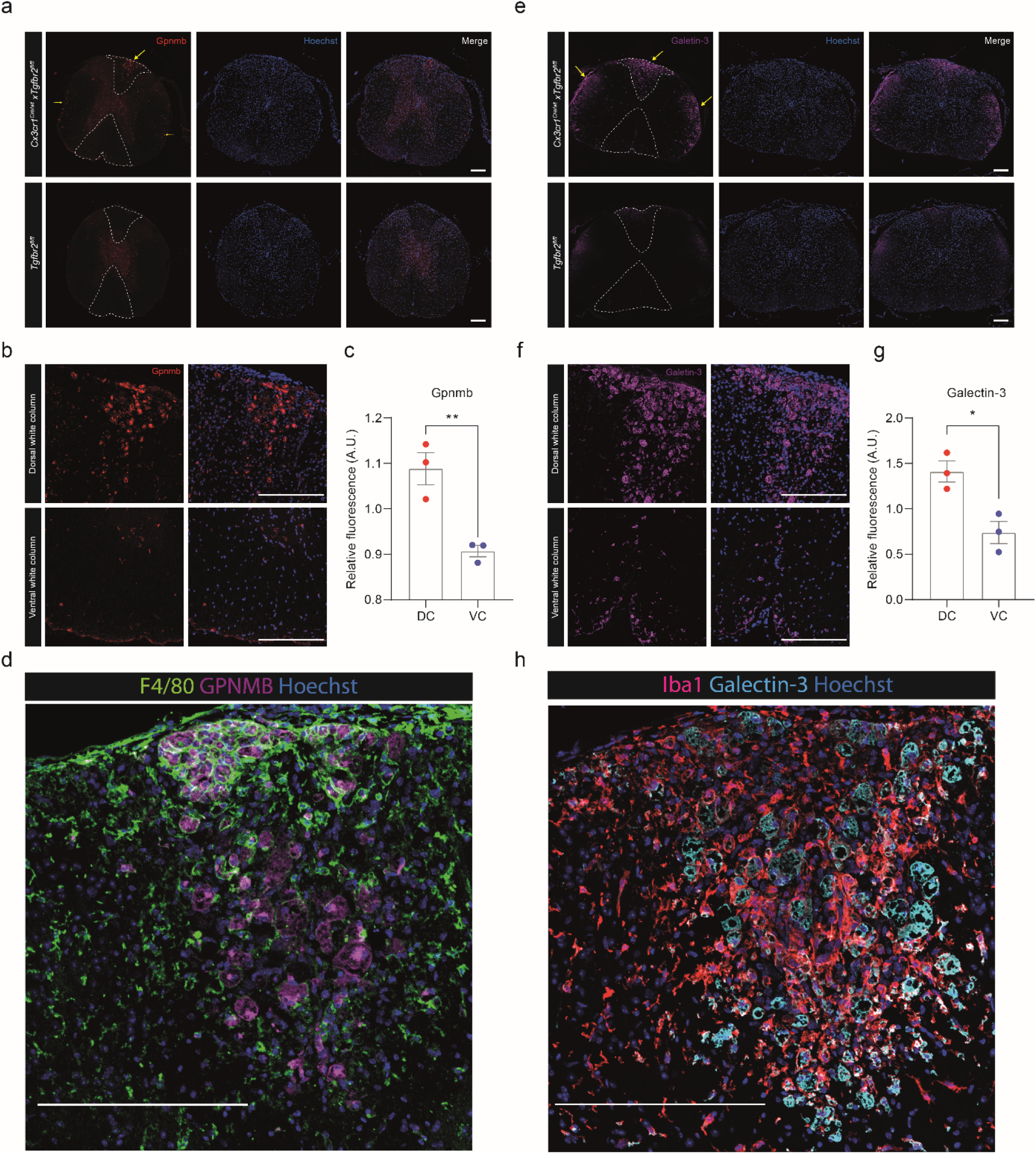
Spatial mapping of GPNMB and Galectin-3 (encoded by Lgals3) expression in the spinal cord. **a**, Representative Immunohistostaining images showing the expression of GPNMB (in red) in the spinal cord of 12-month-old female WT mice and Cx3cr1^CreER^:Tgfbr2 ^fl/fl^ (D30) mice. **b**, Magnification of the circled dorsal column (DC) and ventral column (VC) white matter regions of the Cx3cr1^CreER^:Tgfbr2 ^fl/fl^ mouse in (a). **c**, Quantification of the ratio of GPNMB fluorescent intensity between DC and VC regions within each mouse; n = 3 per group. **d**, Representative Immunohistostaining images showing the expression of GPNMB (in magenta) and F4/80 (in green) in the DC of Cx3cr1^CreER^:Tgfbr2 ^fl/fl^ (D30) mice. e, Representative Immunohistostaining images showing the expression of Galectin-3 (in magenta) in the spinal cord of WT mice and Cx3cr1^CreER^:Tgfbr2 ^fl/fl^ (D30) mice. **f**, Magnification of the circled DC and VC regions of the Cx3cr1^CreER^:Tgfbr2 ^fl/fl^ mouse in (e). **g**, Quantification of the ratio of Galectin-3 fluorescent intensity between DC and VC regions within each mouse; n = 3 per group. **h**, Representative Immunohistostaining images showing the expression of Galectin-3 (in cyan) and Iba1 (in red) in the DC of Cx3cr1^CreER^:Tgfbr2 ^fl/fl^ (D30) mice. Scale bar, 200 μm. Data are shown as mean ± SEM. \**p*⍰<⍰0.05; \*\**p*⍰<⍰0.01; two-tailed Student’s t test.

### Deletion of microglial *Tgfbr2* leads to behavioral abnormalities with gender and age differences

It has been reported that blocking TGF-β signaling impairs microglial stepwise development and results in neurodevelopmental disorders with neuromotor deficits^5,14^. However, disruption of TGF-β signaling in adult microglia does not seem to affect the neurobehavior of adult mice^3^. Notably, only young male mice were used in these studies. Here we included female and male *Cx3cr1^CreER^Tgfbr2^fl/fl^* mice of varying ages (young mice were 8–10 weeks old and older mice were 7–11 months old) and examined the behavior of mice from different aspects, including weight change, tail function, gait assessment, urinary function, limb function, fur assessment, and breathing assessment (Fig. 6 a). We found that around 30 days after microglial *Tgfbr2* deletion, a phase after the onset of DC demyelination, mice began to show aberrant clinical symptoms. Intriguingly, female mice developed more progressive clinical symptoms as compared to the males, and older male mice behaved worse than the young males (Fig. 6 a). We also employed a four-paw hanging wire behavioral test to assess the paw strength of limbs and coordinative function. While the wild-type mice can all stay on the wire until the end (cut-off time: 3 min), microglial *Tgfbr2* deficient female and older male mice start to fall from the wire after 2 weeks of tamoxifen administration; older female mice performed the worst whereas young male mice were only partially affected after 6 weeks of tamoxifen administration (Fig. 6 b). We also kept some microglial *Tgfbr2*- deficient mice for a longer term (76 days after tamoxifen administration) and found that more than half of the older females died or had to be euthanized whereas young male mice all survived (Fig. 6 c). To further limit the effect of *Tgfbr2* deletion on microglia, rather than other CNS macrophages, we also introduced *Tmem119^CreERT2^Tgfbr2^fl/fl^* mice, and similar behavioral phenotypes were observed (data not included yet).

**Fig 6.**
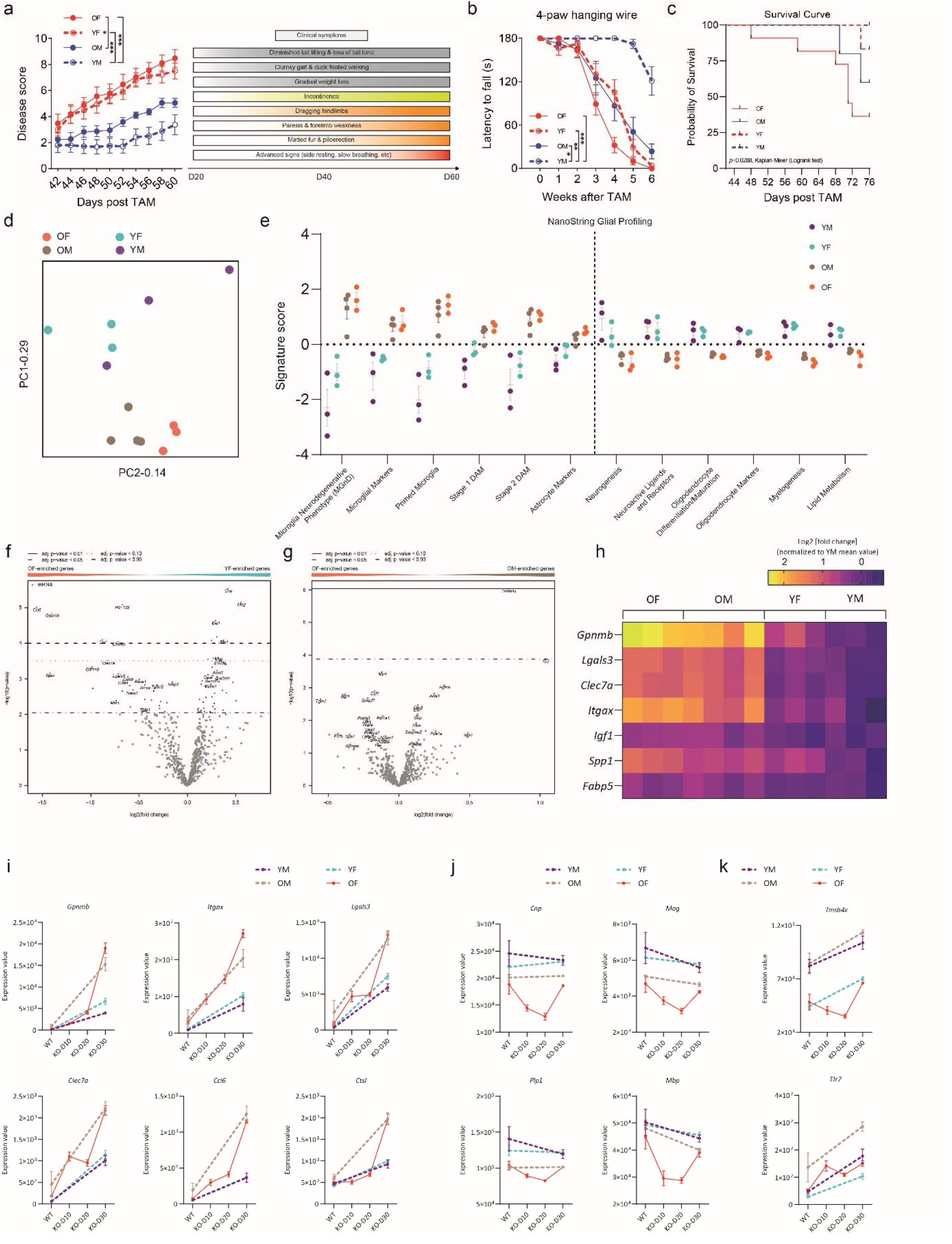
Microglial *Tgfbr2* deficient mice develop severe behavioral deficits with age and gender differences. **a, b** The clinical symptoms of young male (YM), young female (YF), older male (OM), and older female (OF) Cx3cr1^CreER^:Tgfbr2 ^fl/fl^ mice were monitored and scored overtime (young mice were 8- 10 weeks old and older mice were 7-11 months old). A summary of typical clinical symptoms following microglial *Tgfbr2* depletion is presented on the right. Grey boxes indicate early clinical signs starting from around D25 to D30. Yellow-orange boxes indicate aggressed clinical symptoms developed from around Day 30 to Day 40. Orange-red boxes indicate severe, end-stage clinical signs usually seen after Day 40. n = 18 for OF and YF groups; n = 11 and 12 for OM and YM, respectively. **b**, 4-paw hanging wire behavioral tests were performed once a week and the time (cut-off 3 min) mice managed to grasp the wiring lid were recorded. n = 18 for OF and YF groups; n = 11 and 12 for OM and YM, respectively. c, The survival curve of mice up to 76 days following microglial *Tgfbr2* deletion. n = 6 for YM and YF groups; n = 11 and 5 for OF and OM, respectively. **d**, Spinal cord homogenates from YM, YF, OM, OF mice (n = 3-4 per group) after 30 days of microglial *Tgfbr2* depletion were analyzed using the NanoString Glial Profiling panel and the principle component (PC) analysis reveals the separation of the samples/groups. Young mice are 3-4 months old, and old mice are 14-17 months old. **e**, Pathway scores of selected pathways with high relevance to glial cell function from NanoString analysis. **f**, Volcano plot showing the distribution of the differentially expressed genes from NanoString analysis between the YF and OF. Genes to the left are enriched in OF whereas genes to the right are enriched in YF. **g**, Volcano plot showing the distribution of the differentially expressed genes from NanoString analysis between the OM and OF. Genes to the left are enriched in OF whereas genes to the right are enriched in OM. **h**, Heatmap showing the expression of the PAM signature genes across different groups. i, Kinetic expression of selected genes that are enriched in old mice versus young mice following microglial *Tgfbr2* deletion. **j**, Kinetic expression of myelin genes overtime. **k**, Kinetic expression of genes that are enriched in males compared to females in the spinal cord. For (a) and (b), * *p* < 0.05, ** *p* < 0.01, *** *p* < 0.001; area under the curve (AUC) of each group was compared using one-way ANOVA with Tukey’s Multiple Comparison Test. Survival curve in (c) was analyzed using Kaplan-Meier survival analysis with Logrank test for trend.

To understand the molecular mechanisms underpinning the gender and age differences following the loss of microglial *Tgfbr2*, we performed transcriptomic analyses of the spinal cords from the male and female mice at different ages using the NanoString Glial Profiling Panel. PC analyses revealed marked segregation between the young mice and older mice and separation between the older male and female mice, while the separation between the young males and females was not distinct (Fig. 6 d). Pathway analyses revealed that, compared to the young mice, older mice were more enriched in MGnD/DAM signatures, primed microglia signature, and astrocyte/microglial markers, and less enriched in pathways such as *Neurogenesis, Neuroactive Ligands and Receptors, Oligodendrocyte Differentiation/Maturation, Oligodendrocyte Markers, Myelogenesis, and Lipid Metabolism* (Fig. 6 e). To better decipher the age- and gender-associated genes, we did pair-wise comparisons between two groups and demonstrated the DEGs in volcano plots (Fig. 6 f, g). Compared to the young females, the older females downregulated myelin genes such as *Cnp, Mog, Plp1*, and *Mbp* (Fig. 6 f, j), and were enriched in TSM signature genes (*Gpnmb* and *Lgals3*), *Msr1, Apoe, Trem2, Clec7a, Itgax, Ccl6*, and *Ctsl* (Fig. 6 f, i). Many of these genes were also found differentially expressed as core DEGs between the WT microglia and *Tgfbr2*-deleted microglia from our bulk RNA-seq analyses (Fig. 3 b-d). Interestingly, we also noted an increased expression of genes involved in the TGF-β signaling pathway (*Tgfb1, Tgfbr2*, and *Nrros*) in the spinal cords of older female mice as compared to the young females. The signature genes of PAM were also highly induced in the older mice compared to the young mice, further indicating their correlation with the disease (Fig. 6 h). The only DEG we found between older females and males was *Tmsb4x*, which was highly enriched in males (Fig.6 g, k). Of note, *Tmsb4x* encodes thymosin beta-4 and thymosin beta-4 is a potent anti-inflammatory and pro-regenerative peptide and has proven effects on oligodendrocyte differentiation and myelin regeneration^48–54^. The concentration of thymosin β4 in the cerebrospinal fluid of patients with MS is notably reduced compared to individuals with non-MS neurological disorders^55^. However, it should be noted that *Tmsb4x* is an X chromosome gene, and its differential expression between males and females needs to be interpreted with caution regarding its biological relevance to demyelination or remyelination. These data suggest that loss of microglial *Tgfbr2* in adulthood leads to behavioral abnormalities with preference in older and female mice.

### The demand for TGFβ1 in the spinal cord increases with aging under steady state

TGF-β signaling is the goalkeeper for the maintenance of tissue homeostasis^56–58^. The fact that certain areas in the spinal cords are more vulnerable to the loss of microglial *Tgfbr2* and that older mice develop worsened disease prompts us to hypothesize that to maintain tissue homeostasis, the threshold demand for initiating TGF-β signaling might be higher in the DC during aging. Our NanoString analyses suggested that under steady state, the spinal cords of naïve older mice have fewer activities in myelin genesis and oligodendrocytes differentiation, but more MGnD and cytokine activities, as compared to that of the naïve young mice (Fig. 7 a). Interestingly, we found that the mRNA expression of *Tgfb1* increased with age in female spinal cords, and a similar trend was observed in the males (Fig. 7 b). Furthermore, the expression of *Tgfbr1* and *Tgfbr3* was also found to be upregulated in older females compared to young females (Fig. 7 b). We further confirmed an increased protein expression of TGFβ1 and *Tgfbr2* with aging in the spinal cords by Western blotting, but SMAD2 expression and its phosphorylation remained unchanged, indicating that more TGFβ1-TGFBR2 were needed during the aging process to support sufficient transduction of TGF-β signaling in the spinal cords (Fig. 7 c, d). Interestingly, the ratio between latent TGFβ complex and dimeric TGFβ also increased with aging, suggesting the extracellular liberation of bioactive TGFβ from its latent TGFβ complex may also decrease in older mice (Fig. 7 d). To further characterize subregional differences in the expression of genes involved in the TGF-β signaling pathway, we micro-dissected the DC, VC, and GM regions of naïve mice at young or older ages and compared the mRNA expression of these genes (Fig.7 e). In general, compared to the GM, the expression of the genes in the TGF-β signaling pathway was higher in the DC and VC, and there was also an increasing trend with aging. Of note, we found elevated expression of *Tgfb1, Tgfbr1*, and *Tgfbr2* in the DC and VC in older mice compared to the young mice, but their expression remained unaltered with aging in the GM. In the DC of older mice, the expression of *Tgfb1, *Tgfbr2*, Tgfbr3*, and *Smad4* was significantly higher than that in the VC and GM (Fig. 7 e). Meanwhile, the expression of genes that are downstream negative regulators of the TGF-β signaling, including *Smad7, Smurf1*, and *Smurf2*, was also significantly elevated in the DC of older mice compared to that in the VC and GM (Fig. 7 e). This suggests that in the DC of older mice, the efficiency of transducing upstream TGFβ1-TGFBR2 binding signals to induce the expression of TGF-β target genes may also be impeded. Therefore, DC may demand more TGFβ1 to maintain a functional net effect. This was further confirmed by immunostaining showing a higher abundance of TGFβ1 in the DC of naïve older mice as compared to the VC (Fig. 7 f, g). We also noted that in the VC, TGFβ1 was mainly located inside the IBA1+ microglia, whereas in the DC, TGFβ1 was not only colocalized with microglia but was also expressed in the surrounding areas of microglia (Fig.7 g). The observation of TGFβ1 expression outside of microglia in the DC suggests that microglia in this region may be exposed to a TGFβ1-rich environment under steady state conditions. Taken together, these findings suggest that there is an increased demand for TGFβ1 in the spinal cord with aging, particularly in the DC, which could explain why the disruption of TGF-β signaling in microglia leads to demyelinating pathology in the DC and worsened disease in older mice.

**Fig 7.**
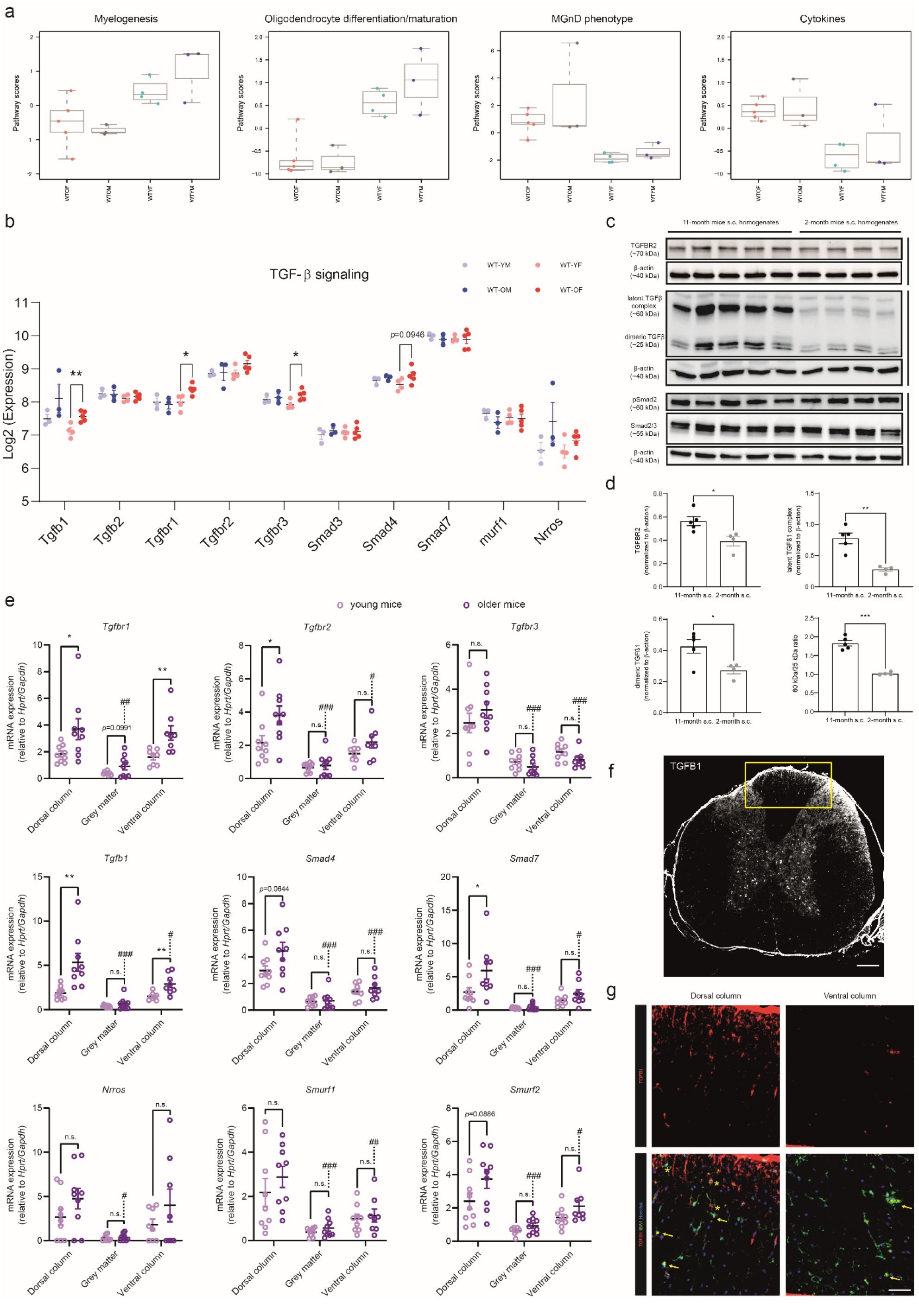
Characterization of TGF-β signaling in the spinal cord. **a**, Pathway score of selected pathways from NanoString analysis of the spinal cord homogenates of naïve older female (OF), older male (OM), young female (YF), and young male (YM) mice. **b**, The normalized mRNA expression (Log2-transformed from NanoString results) of genes in the TGF-β signaling pathway across the OF, OM, YF, and YM groups. **c, d,** Western blotting of *Tgfbr2*, TGFB1, pSMAD2, and β-actin expression in the spinal cord homogenates of young (2-month-old) and older (11-month-old) male mice with quantification in (d). n = 4-5 per group. **e**, RT-PCR results showing the mRNA expression of genes in the TGF-β signaling pathway across the DC, VC and GM regions in 8-week-old and 5-8-month-old mice (both male and female mice are included). n = 8-10 per group. **f**, Immunohistostaining showing the overall distribution of TGF-β1 in the spinal cord. Yellow box highlights the enriched TGF-β1 expression in the dorsal column. Scale bar, 200 μm. **g**, Co-staining of Iba1 (red), TGF-β1 (green), and Hoechst (blue) in the DC and VC regions. Yellow arrows indicate microglia with TGF-β1 production, and yellow asterisks indicate microglia surrounded by TGF-β1. Scale bar, 50 μm. Data are shown as mean ± SEM. \**p*⍰<⍰0.05; \*\**p*⍰<⍰0.01; \*\*\**p*⍰<⍰0.001; two-tailed Student’s t test. #*p*⍰<⍰0.05; ##*p*⍰<⍰0.01; ###*p*⍰<⍰0.001; compared to the DC of older mice using one-way ANOVA with Dunnett’s Multiple Comparison Test.

## Discussion

Myelination and the maintenance of myelin are complex processes requiring coordinated cell-cell interactions involving axons, myelin-forming cells, and support from microglia and astrocytes^59,60^. Firstly, the properties of neurons/axons are determinants of myelination^61,62^. Adenosine produced in axons during action potential firing activates the adenosine receptors expressed on OPCs and induces their differentiation^63^. The polysialylated neural cell adhesion molecule (PSA-NCAM) expressed on the surface of axons negatively regulates myelination as it may prevent oligodendrocytes from attaching to the axons^64,65^. The axon-derived neuregulin-1 (NRG1) acts on epidermal growth factor receptors (EGFR), such as ErbB2/3, expressed on myelin-forming cells, representing one of the most well-studied and key signals of myelination and regulation of myelin thickness^66–70^. Secondly, fine-tuned microglia-astrocyte-oligodendrocyte crosstalk is fundamental for oligodendrocyte differentiation and myelin maintenance^71,72^. TGFβ1 released predominantly by microglia in the CNS represents one of the most important cytokines regulating postnatal myelin growth and integrity^73,74^. A recent study firmly demonstrated that microglia maintain myelin health in adult mice and humans, and that disruption of TGFβR1 signaling in oligodendrocytes affects myelin integrity^75^. Cytokines released by microglia and astrocytes under pathological conditions, such as IL-1β, TNF, and IFN-γ, activate and recruit OPCs and facilitate their proliferation^76,77^, but may also lead to the loss of mature oligodendrocytes^29,78–81^. Therefore, a trivial change in any of the cells involved may trigger a butterfly effect and disrupt myelin integrity.

In light of the aforementioned facts, putative theories are proposed in this study regarding why the DC is more vulnerable to demyelination in the absence of microglial TGF-β signaling compared to the VC: i) microglia in the DC are different from their counterparts in the VC, and they respond differently to the loss of TGF-β signaling. ii) the microglial response following *Tgfbr2* deletion is homogeneous, but the myelin or mature oligodendrocytes are heterogenous and those in the DC are more vulnerable to the damage caused by *Tgfbr2*-deficient microglia. iii) the neuronal and astrocytic activities in the DC are more sensitive to microglial dysfunction due to the lack of *Tgfbr2*, and myelin loss in the DC is a secondary response to the deregulated neuronal/astrocytic activities. With these hypotheses, our snRNA-seq data and transcriptomic analyses of sorted microglia from DC, VC, and GM reveal tentative information. The changes in the populations of neurons after microglial *Tgfbr2* deletion appeared to be minor, suggesting that neuronal deregulation on oligodendrocytes may not be the leading cause of the pathology. We noted a slightly downsized cluster of astrocytes which could be a relative decrease in frequency as the microglial cluster expanded; further investigations may be needed to explore astrocyte contribution, and it would be of interest to test whether disruption of TGF-β signaling in astrocytes could also lead to a similar phenotype as observed caused by microglia. By contrast, changes in microglia and oligodendrocyte lineage cells were obvious. However, although we revealed a certain number of DEGs between the DC and VC/GM microglia and a higher level of inflammatory signatures in the DC microglia from our RNA-seq analyses, the overall difference between DC and VC/GM microglia was not considerable, and it is still questionable whether the elevated expression of *Il1b* and *Tnf* in the DC microglia could lead to such a different pathological phenotype, although these cytokine genes were highly relevant to myelin/oligodendrocyte abnormalities^25,78,82–85^. Of note, in the cuprizone-induced rodent demyelinating model, elevated expression of *Il1b, Tnf*, and *Il6* in the corpus callosum, the most vulnerable demyelinating site following cuprizone treatment, precedes demyelination^86^. Future studies administering exogenous IL-1b and TNF to spinal cords to see whether and where spontaneous demyelination occurs in the spinal cords will provide more information regarding their contribution to the DC demyelination as observed in our model. On the other hand, the ontogeny of myelin and oligodendrocyte in the DC and VC is quite different^87^. During development, oligodendrogenesis in the VC of the developing spinal cords is mainly ascribed to the motor neuron progenitor (pMN) domain and is dependent on the *Shh* signaling and the homeodomain transcription factor *Nkx6*^88,89^. By contrast, a wave of late-phase oligodendrogenesis from the dorsal neural tube is independent of Shh and *Nkx6* gene activities, and retains a dorsal identity, suggesting differences in the molecular control of ventral and dorsal oligodendrocytes^90–92^. Surprisingly, the spatial distribution of the dorsally derived oligodendrocytes is mainly located in the dorsal and dorsolateral columns of the spinal cord, which well matches the affected regions in our study^93^. We therefore speculate that disrupting microglial TGF-β signaling may specifically target the dorsally derived oligodendrocytes. Further studies inducing microglial *Tgfbr2* deletion in the fate mapping mouse tool labeling dorsal oligodendrocytes may help prove this hypothesis^93^. It remains unsolved whether the dorsal oligodendrocytes are functionally equivalent to the ventral oligodendrocytes, and whether the damaged myelin and mature oligodendrocytes in the DC are developmentally different from their ventral counterparts, which warrants future investigations. Previous studies reported similar electrical properties of dorsally and ventrally derived oligodendrocytes^93,94^, but oligodendrocyte progenitors with a dorsal origin exhibited enhanced ability in proliferation, migration, and differentiation, and are more susceptible to age-associated differentiation impairment^95^. Therefore, we are inclined to conclude that demyelination in the DC following microglial *Tgfbr2* deletion may result from both heightened proinflammatory activation of microglia in the DC and an increased functional dependence of DC oliogdendrocytes on steady-state TGF-β signaling.

It is also noteworthy that in our previous study, we also found that compared to the VC region, the DC region is more vulnerable to demyelination when the CNS microglia are replaced by *Tgfbr2*-deficient monocyte-derived macrophages. This suggests that the myelin columns in the DC region, rather than the microglia, are imprinted with specific local environmental cues (exemplified by being immersed in a higher TGFβ1 environment under homeostatic states) or bear a different developmental trajectory, which makes them more dependent on fine-tuned TGF-β-mediated cell-cell interactions. In line with this, a recent study also identified a *Klk6*^+^ mature oligodendrocyte subcluster that is spatially specifically enriched in the DC and almost absent in other brain regions, such as corpus callosum and cortex^96^. The ontogeny and function of these *Klk6*^+^ mature oligodendrocytes remain mysterious, and it is yet to be confirmed whether the lost myelin/oligodendrocytes in the DC following microglial *Tgfrb2* deletion are those *Klk6*^+^ oligodendrocytes or dorsally derived oligodendrocytes.

Despite all this, we reveal substantial molecular changes following microglial *Tgfbr2* deletion. Apart from the prominent inflammatory signature, we also noted induced gene sets that are highly relevant to myelin damage, such as the matrix metalloproteinases (MMPs) and cell adhesion molecules (CAMs), including the C-type lectin/C-type lectin-like domain (CTL/CTLD) superfamily. These genes encode proteins that are important regulators of the ECM and mediate cell-cell interactions, and are also crucial extracellular molecules for liberating TGFβ^32,97^. The increased expression of MMPs (*Mmp9, Mmp12*, and *Mmp13*) and CAMs in microglia following *Tgfbr2* deletion, together with an increased level of Tgfb1 in the spinal cord as revealed by our NanoString analyses, indicate that there might be a compensatory reaction to the loss of *Tgfbr2*: other cells in the spinal cords are producing more *Tgfb1*, and meanwhile, more MMPs and ECM molecules are induced by microglia to facilitate local activation of TGFβ on the microglial surface. These response in microglia may endow them with a ‘gluey’ property, strengthening their attachment to myelin and may physically disturb them^98,99^; abundant MMPs produced by *Tgfbr2*- deleted microglia could further degrade myelin components^33,100–105^, thus leading to demyelination. It is noteworthy that TSMs, induced following *Tgfbr2* deletion as revealed by our snRNA-seq analyses, strongly expresses *Mmp12*, which has been widely recognized for its damaging effects on myelin integrity, highlighting the pathogenic role of TSMs^106–110^. Other notable genes of TSMs include *Lgals3, Gpnmb*, and *Mgll*. Growing evidence supports a proinflammatory role of Lgals3 in microglia and neuroinflammatory conditions^111–113^, as well as ingesting myelin^114^, and Gpnmb has been identified as a significant risk factor for neurodegenerative diseases^115,116^. Additionally, the foamy cells in the rim of chronic active MS plaques highly express GPNMB, and it is also highly induced in microglia/macrophages outside the rim of lesions^117^, suggesting an active role of GPNMB in lesion expansion. There is limited information available on the function of *Mgll* on microglia, but *Mgll* encodes monoacylglycerol lipase, which disrupts myelin integrity^118,119^. Further research is needed to understand how the microglia-derived *Gpnmb, Lgals3*, and *Mgll* affect other cell types, particularly oligodendrocytes, and this may lead to the identification of potential therapeutic targets for demyelinating diseases.

Our findings demonstrated that the elimination of *Tgfbr2* in adult microglia has an impact on microglia-specific gene signatures, contrary to previous findings that such signatures would remain unchanged upon disruption of the TGF-β signaling pathway ^3,4^. This discrepancy could be a result of the different ages and gender of mice used for microglial transcriptomic analyses. In addition, in the previous study using the same *Cx3cr1^CreER^Tgfbr2^fl/fl^* mice, behavioral deficits following microglial deficiency of TGF-β signaling in adult mice were not revealed, whereas in our study, we observed severe demyelinating disease with neurobehavioral dysfunction, and there is a clear gender difference that recedes with aging. When comparing our study with the previous one in detail, we found that only young male mice (∽8 weeks) were used in the previous study. In this regard, it is consistent with our finding showing that young male mice were only mildly affected by the loss of microglial TGF-β signaling, whereas female and older mice were much more vulnerable to developing clinical symptoms and behavioral deficits. This underscores the heterogeneity of microglia under the influence of gender-associated and age-associated factors, and thus it is important to take into consideration gender and age when conducting microglia-related studies, as stated before also by others ^120–124^.

It is interesting to find that the expression of TGFβ1-TGFBR2 in the spinal cords increases with aging, which suggests that the threshold for upstream input of the TGF-β signaling in older mice to maintain a homeostatic state is higher. This could also explain why older mice are more prone to develop behavioral deficits following the disruption of TGF-β signaling. However, deleting *Tgfbr2* in microglia does not seem to affect microglial *Tgfb1* expression, and the DC region or DC microglia appear to bear the highest *Tgfb1* expression under homeostatic states or after *Tgfbr2* deletion. This implies that the DC regions or DC microglia may require more TGF-β signaling to maintain homeostasis and therefore are more sensitive to the deletion of *Tgfbr2*. In addition, the downstream execution of TGFβ1-induced signaling transduction in the DC may not be efficient, especially in older mice, as evidenced by increased expression of inhibiting genes (*Smad7, Smurf1, Smurf2*) of the TGF-β signaling in the DC, and more TGFβ1 is thus demanded to maintain a similar expression level of the target genes of the TGF-β signaling under homeostatic states. Detailed characterization would be needed to provide more solid evidence, and it would add more value if similar findings could be recapitulated in human spinal cords. Human neurological diseases such as copper deficiency myelopathy and subacute combined degeneration primarily exhibit degenerative pathologies in the dorsal column of the spinal cord^125–129^, with accumulation of foamy microglia^130,131^. It would be of interest to investigate the involvement of dysregulated microglial TGF-β signaling in these human diseases.

There are limitations in our study that could be dispelled by further investigations. We have revealed *Tmsb4x* as a potential protective factor for male mice, and although several studies have confirmed its role in facilitating oligodendrocyte differentiation and remyelination, further studies will be needed to confirm whether giving thymosin beta-4 (encoded by *Tmsb4x*) to female and old mice in microglia *Tgfbr2*-deficient mice could rescue their disease progression. Similarly, loss-of-function studies downregulating microglial *Gpnmb, Lgals3*, and *Mgll*, such as using microglia-specific targeting tools as we previous developed^132^, would further confirm their function and contribution to the demyelinating pathology observed in our study. In addition, the age-dependent and region-specific TGFβ expression pattern in the spinal cords shall be validated in human settings.

In conclusion, our study provides new insights into the regulation of adult microglial homeostasis by TGF-β signaling and its impact on myelin integrity and the development of demyelinating diseases. These findings substantiate our understanding of the complex interplay between microglia and the central nervous system and the potential for targeting microglia as a therapeutic strategy for demyelinating disorders.

## Methods

### Experimental subjects

Experimental mice were bred and housed at the Comparative Medicine Department at Karolinska University Hospital, Sweden. Mice were maintained in climate-controlled and pathogen-free environment with regulated 12-h light/dark cycles. *Cx3cr1^CreER-YFP^* (#021160), *Tmem119^CreER^* (#031820), *Ccr2^CreE-GFP^* (#035229) and *Tgfb1 ^fl/fl^* (#065809) mice were obtained from the Jackson Laboratory. *Tgfbr2*^fl/fl^ mice were kindly offered by M. Li at Sloan Kettering Institute. C57BL/6NTac mice (Taconic) were bred at the local animal facility. For tamoxifen administration, tamoxifen (T5648, Sigma) was dissolved in corn oil at 75 °C for 1 h; mice were administered three injections of 4– 5 mg tamoxifen in 200 μL corn oil (i.p) with a one-day interval between each injection. The day mice received the last tamoxifen injection was counted as Day 0 (D0). Both male and female mice were included for most experiments, and generally experimental mice were 2-4 months old unless specified for age-related comparisons: young mice were around 2 months old, and older mice were around 7-12 months old. For bulk RNA-seq, female mice aged around 4 months were used; for snRNA-seq female mice aged around 12 months were used. Mice were euthanized by injection of 100 μL of pentobarbital sodium (i.p), and perfused with cold HBSS.

### Ethics statement

Animal experiments were performed under the guidelines from the European Community Council Directive and the Swedish National Board for Laboratory Animals (86/609/EEC) under the ethical permits N138/14, followed by 9328-2019, and 23561-2022, approved by the local ethical review board the North Stockholm Animal Ethics Committee.

### Immunofluorescence

Mouse spinal cord tissues were fixed with 4% PFA for 24⍰h followed by dehydration in 20 and 30% sucrose at 4 °C. The dehydrated tissues were embedded with OCT cryo-mountant (45830, Histolab) and kept at −80 °C. OCT tissue chunks were sectioned (14 μm) in a Leica CM1850 cryostat and mounted on SuperFrost Plus Adhesion slides (J1800AMNZ, Epredia) kept at −20°C. A mild antigen retrieval step was introduced by immersing the slides into boiled antigen retrieval solution (00-4955-58, Invitrogen) for 10 min, followed by washing with PBS and incubation with blocking buffer (5% normal goat- or donkey-serum and 0.3 % Triton X-100 in PBS) for 1 h at room temperature. The following primary antibody were used: anti-IBA1 (1:500, 019-19741, WAKO), anti-MHC-II (1:200, 46-5321-80, Thermo), anti-F4/80 (1:200, MCA497GA, Bio-Rad), anti-GPNMB (1:500, ab188222, Abcam), anti-Galectin-3 (1:100, 14-5301-82, Thermo), anti-TGFB1 (1:500, MA1-21595, Thermo). Primary antibodies were diluted in staining buffer (1% BSA and 0.3 % Triton X-100 in PBS) and incubated for overnight at 4 °C, followed by washing and staining with species-matched secondary antibodies for 1 h at room temperature. For Fluoromyelin (1:300, F34651, Invitrogen) staining, sections were incubated for 30 min at room temperature. Nuclei staining were performed by incubating with Hoechst (62249, Thermo). Images were taken using a Zeiss LSM880 confocal microscope and images were analyzed using the Zeiss ZEN software or FIJI/ImageJ.

### Microdissection of spinal cord tissue

After perfusion and collection of the spinal cords, they were either immersed in cold HBSS and microdissected right after for flow cytometry and cell sorting or immersed in RNAprotect Tissue Reagent (76104, Qiagen) for mRNA analyses. Microdissection was performed under a Leica MZ95 surgical microscope. The DC was gently pulled out using ophthalmic forceps curved with hook, and the VC was carefully dissociated using iris scissors and scalpels; GM was carefully separated with connecting white matter tissues and clapped out using iris scissors and tweezers.

### Preparation of CNS single-cell suspensions

After perfusion, spinal cord tissue was flushed out from the spine, and physically dissociated followed by enzymatic digestion with papain (1:100 diluted in L15 medium; LS003126, Worthington) and 0.2 mg/mL DNase I (10104159001, Roche). Myelin was removed using 38% isotonic Percoll (P1644, Sigma) by centrifugating at 800 g (4x acceleration, no brake) for 10 min. Cell pellets were suspended for flow cytometry or cell sorting.

### Flow cytometry

Spinal cord single cell suspensions were stained with LIVE/DEAD Fixable Stain Kits (1:1000, L10119 or L34959, Invitrogen) to exclude dead cells. The following antibodies were used: PerCP-Cy5.5 anti-CD11b (1:100, 101228, BioLegend), PE-Cy7 anti-CD45 (1:100, 103114, BioLegend), PE anti-Ly6C (1:250, 128008, BioLegend), APC anti-F4/80 (1:100, 123118, BioLegend), A700 anti-MHC-II (1:100, 107622, BioLegend).

### Cell sorting

Single cell suspensions from the microdissected spinal cord subregions (DC/VC/GM) were sorted using a BD Influx cell sorter. After myelin removal, the single cell suspensions were stained with LIVE/DEAD Fixable Yellow Dead Cell Stain kit (1:1000, L34959, Invitrogen), and surface antibodies: PerCP-Cy5.5 anti-CD11b (1:100, 101228, BioLegend), PE-Cy7 anti-CD45 (1:100, 103114, BioLegend), PE anti-Ly6C (1:250, 128008, BioLegend), and APC anti-F4/80 (1:100, 123118, BioLegend). Microglia cell were sorted as CD11b^+^CD45^+(int)^Ly6C^-^F4/80^+(int)^; cell purity was determined to be > 95% by checking the YFP expression of the gated *Cx3cr1^CreER-YFP^* cells. A total of 100 microglia from each region were collected in 5 μL single-cell lysis buffer (635013, Takara), and flash-frozen on dry ice.

### Next-generation sequencing

The sorted 100 cells from each spinal cord subregion were proceeded directly with cDNA synthesis using the SMART-Seq v4 Ultra Low Input RNA Kit, and library was prepared using the Nextera XT kit by staff at the Bioinformatics and Expression Analysis (BEA) core facility at Karolinska Institutet. For each group, N = 3–5 biological samples were included. Next-generation sequencing was performed at the National Genomics Infrastructure (NGI) at the Science for Life Laboratory using a NovaSeq 6000 S4 platform. Data normalization was performed using the DESeq2 method^133^; differential gene expression and KEGG_Pathway/GO Enrichment analysis were performed using the Dr. Tom bioinformatic analysis tool (BGI).

### Nuclei isolation and snRNA-seq

Nuclei isolation was performed following a 10X Genomics protocol (CG000124) with modifications. Nuclei lysis buffer were freshly prepared: 10 mM Tris-HCL (T2194, Sigma), 10 mM NaCL (59222C, Sigma), 3 mM MgCL_2_ (M1028, Sigma), 0.1% Nonidet P40 substitutet (74385, Sigma) in PBS. Spinal cord tissue (1- cm long segment) was immersed in 2 mL lysis buffer and homogenized using a dounce homogenizer on ice until most nuclei were released when checking under a microscope. Homogenates were added with 5 mL Hibernate medium and passed through a 30 μm cell strainer, followed by centrifuging at 500 g for 5 min at 4 °C. The pellets were washed twice by being resuspended in 10 mL washing buffer (2% BSA and 1 mM DTT in PBS supplemented with RNase inhibitor) followed by centrifugation as previous settings. After washing, the pellet was resuspended in 38% isotonic Percoll (P1644, Sigma) and centrifuged at 2000 *g* (4x acceleration, no brake) for 20 min at 4 °C. The floating myelin layer was carefully removed, and the pellets were immediately resuspended in wash buffer in low-binding Eppendorf tubes. For each biological group/timepoint, spinal cord segments from two mice were pooled. Nuclei were counted and a final concentration of around 1000 nuclei/μL was used. Library was constructed using the Chromium Next GEM Single Cell 31 Reagent Kits v3.1 (Dual Index; CG000315 Rev), QC was performed using Bioanalyzer (Agilent), concentration was determined using Qubit (Thermo Fisher) and KAPA Library Quantification Kits (Roche) following the manufacturer’s protocol. Samples were sequenced on NovaSeq6000 (NovaSeq Control Software 1.7.5/RTA v3.4.4) with a 28nt(Read1)- 10nt(Index1)-10nt(Index2)-90nt(Read2) setup using ‘NovaSeqStandard’ workflow in ‘SP’ mode flowcell. The Bcl to FastQ conversion was performed using bcl2fastq_v2.20.0.422 from the CASAVA software suite. The quality scale used is Sanger / phred33 / Illumina 1.8+.

### snRNA-seq preprocessing and unsupervised clustering

SnRNA-seq data for each sample was preprocessed using the cellranger count command in the Cell Ranger toolkit (version 7.0.0) provided by 10x Genomics for the alignment against the mouse reference genome (mm10), followed by filtering, barcode counting, and unique molecular identifier (UMI) counting to generate gene-barcode matrices. The Seurat package (version 4.2.1) was subsequently used to generate the gene expression matrix. For quality control, we kept high quality cells with thresholds of 800-3000 unique gene counts, and less than 20% mitochondrial counts. After quality control, the Seurat object of each sample was normalized using the SCTransform command and integrated based on the 5000 most variable genes for dimensional reduction and clustering. Principal component analysis (PCA) was performed on the gene expression matrix and the shared nearest neighbor (SNN) graph was constructed using the FindNeighbors command based on the top 50 principal components. The Louvian algorithm was applied on such nearest neighbor graph to find cell clusters using the FindClusters command with the resolution parameter of 0.3. The same principal components were used for dimensional reduction to generate the Uniform Manifold Approximation and Projection (UMAP) for visualization.

### Identification of cell types

Clusters of different cell types were manually identified based on the expression of signature genes. Briefly, the FindConservedMarkers command in the Seurat package was used to identify differentially expressed genes for each cluster and the cell type annotation for each cluster was confirmed by elevated expression of known cell type signature genes (Supplementary Fig. 2 a): MG for microglia (*Runx1, Ctss, Ptprc, Inpp5d, Mertk*), ASTRO for astrocytes (*Slc39a12, Gli3, Aqp4*); OPC-1 (*Pdgfra, Cspg4, Tnr*), OPC-2 (*Tcf7l2, Bcas1, Sox6, Tnr, Mbp*), OL-1 (*Prr5l, St18, Gab1, Rnf220, Mog, Mbp, Plp1, Mag*), OL-2 (Prr5l, Mobp, Mag, Mbp, Plp1, Mog), and Myelin_OL (*Plp1, Mag, Prr5l, St18, Mobp, Mog*) for cells of the oligodendrocyte lineage; PC for pericytes (*Vtn, Rgs5, Abcc9, Pdgfrb, Flt1*); MN for motor neurons (Chat, Pdgfd, Slit3, Pappa), and PGC for preganglionic motor neurons (*Nos1, Chat*). For the other neuron cell types, we referred to the SeqSeek open-source platform for annotation^39^. Briefly, *Slit2, Nrg1*, and *Robo1* were used as pan-neurons marker, with a combination of markers such as *Cpne4, Reln, Sox5, Adarb2, Ebf2, Nmu, Pde11a, Hmga2, Chst9, and Ptprk used for identifying excitatory neuron (ExcitNeu) clusters, and Rorb, Stac, Cdh3, Gad1, Pdzd2, Adamts5, Gpc5, Zmat4, Nxph1, Rxfp2*, and *Nfib* used for inhibitory neuron (InhibNeu) clusters.

### Analysis of microglia cells

Identified microglia were extracted for the subsequent analyses. PCA was first performed on the gene expression matrix of extracted cells and the SNN graph was constructed with the top 10 principal components. The Louvian algorithm was applied on such nearest neighbor graph to find microglia subclusters with the resolution parameter of 0.4. The same principal components were used for dimensional reduction to generate the UMAP for visualization. The differentially expressed gene analyses between different microglia subclusters were performed with the FindMarkers command in the Seurat package. Gene set enrichment analysis (GSEA) based on gene ontology (GO) was conducted using the clusterProfiler package (version 4.4.4) with a significant cut-off of adjusted P value < 0.05. The monocle package (version 2.24.1) was used for pseudotime trajectory analysis of extracted microglia cells. Differentially expressed genes among three microglia subclusters were identified using the differentialGeneTest command. Dimensional reduction, cell ordering and pseudotime trajectory construction were performed on these differentially expressed genes with default parameters. Branched expression analysis modeling (BEAM) in the monocle package was applied to identify genes with branch-dependent expression.

### NanoString nCounter Glial Profiling

The total RNA of the spinal cords was extracted using TRIzol Reagent (A33250, Invitrogen) following the manufacturer’s protocol. For each group, n = 3-5 biological samples were included. The concentration of the total RNA was measured using a Qubit RNA HS Assay kit (Q32852, Invitrogen), and samples were adjusted to RNA concentration of 20 ng/μL before sending to KIGene Core Facility at Karolinska Institutet for analysis. The samples were first hybridized to the nCounter Mouse Glial Profiling Panel CodeSet (XT-CSO-M-GLIAL-12, NanoString). The CodeSet includes 757 probes for genes of interest, 13 probes for internal reference genes, and 10 additional customized probes for *Smad4, Smad7, Nrros, *Tgfbr2*, Tgfbr3, Smurf1, Tgfb2, Cxcr4, Adgre4*, and *Clec4b1*. All the probes were single target-specific color-coded probes with around 100 bases in length and no reverse transcription or amplification process were needed. After hybridization, the target-probe mix were purified using an automated fluidic handing nCounter Prep Station and excessive probes were eliminated. The target-paired probes were subsequently immobilized in a sample cartridge, and the fluorescent reporter probes were analyzed using the nCounter Digital Analyzer. The mRNA reads were normalized with the 13 internal reference genes included in the CodeSet; data normalization and further advanced analyses were performed using the NanoString nSolver Analysis Software for the provided guidelines.

### Clinical scoring

Clinical symptoms were examined after tamoxifen administration. We assessed weight loss, tail tonus, tail lifting, gait pattern, dragging of legs, incontinence, paralysis, fur abnormality, and breathing abnormality. Mice were assigned one point for the presence of each of the indicated symptom. Mice that reached humane endpoint or died were scored 15 from the day they were euthanized or died.

### Hanging-wire test

Four-paw hanging-wire test was performed once per week. Mice were placed on a wire cage lid and inverted over a cage with soft beddings. The time until each mouse fell from the lid was recorded. The cut-off time was set to 180 s and normal mice could all reach this time. Two trials were performed for each mouse if the mouse did not stay until 180 s during the first trial, and the longest time was recorded.

### Quantitative reverse transcription-polymerase chain reaction (qRT-PCR)

Microdissected DC, VC and GM were processed using a RNeasy Lipid Tissue Mini Kit (74804, Qiagen) to extract total RNA with on-column DNase digestion using a RNase-Free DNase Set (79254, Qiagen). cDNA was synthesized using an iScript kit (1708891, Bio-Rad), and qRT-PCR was performed with SYBR green reactions (1708886, Bio-Rad) in a 384-well plate run in a Bio-Rad CFX384 Touch Real-Time PCR Detection System.

### Western blotting

Spinal cords were homogenized and sonicated in RIPA buffer (R0278, Sigma) supplemented with protease/phosphatase inhibitors (1861261, Thermo) and phenylmethylsulfonyl fluoride (36978, Thermo). Homogenates were centrifuged (11,000⍰g for 20 min at 4°C) and the supernatants were determined using a BCA protein assay kit (23225, Thermo). Samples with equal amounts of proteins were added to 4× Laemmli sample buffer (1610747, Bio-Rad) supplemented with DTT (10197777001, Sigma), and electrophoresed in 4–20% Mini Protean TGX precast protein gels (4561093, Bio-Rad), followed by transferring to a 45⍰mm nitrocellulose membrane (1620115, Bio-Rad). After blocking with 5% non-fat dried milk in TBS-T (20⍰mM Tris-HCl and 137⍰mM NaCl with 0.1% Tween 20 at pH 7.6) for 30⍰min at room temperature, the membrane was incubated overnight at +4°C with the following primary antibodies: anti-TGFBR2 (1:1000, SAB5700739, Sigma), anti-TGFB1 (1:1000, MA121595, Thermo), anti-SMAD2/3 (1:1000, 8685T, CST), and anti-phospho SMAD2 (1:1000, AB3849-I, MERCK). After rinsing, the blots were incubated with HRP-linked secondary antibodies for 2⍰h at room temperature. The blots were incubated with Clarity Western ECL (170-5061, Bio-Rad) and chemiluminescence was detected using a Bio-Rad Chemidoc Imaging System. Band intensity was quantified using the Bio-Rad Image Lab software, and data were normalized to β-actin as the reference protein.

### Statistical analyses

Analyses and graphs were performed using GraphPad Prism 9.5.1 software. Comparison between two groups was done with Student’s two-tailed unpaired t test. Multiple comparison test with a control group was done with one-way ANOVA and Dunnett’s multiple comparison test. Multiple comparison test comparing the mean with every other group was done with one-way ANOVA and Tukey’s multiple comparison test. *p*⍰<⍰0.05 was considered statistically significant. Analysis of RNA-seq data was done with the DESeq2 package using Wald testing with Benjamini–Hochberg adjustment for multiple testing. Detailed statistical information was provided in the figure legends.

## Acknowledgements

We thank the staff at our animal facility (AKM) for their animal caretaking, X. Li (KI) for help with nuclei isolation and single cell data analysis. We thank J. Han, X.M. Zhang, I. Benito-Cuesta, M. Khademi and S. Lewandowski at KI for help with experimental materials, lab management, and scientific discussions. We thank A. van Vollenhoven (KI) for cell sorting.We thank KIGene staff (K. Gell, A. Eriksson, M. Alvehus) and J. Zhang for help with NanoString setup. We thank F. Fagerström, S. Fält, and D. Brodin at BEA (KI) for RNA-seq setup. We acknowledge support from the National Genomics Infrastructure in Stockholm funded by Science for Life Laboratory, the Knut and Alice Wallenberg Foundation and the Swedish Research Council, and SNIC/Uppsala Multidisciplinary Center for Advanced Computational Science for assistance with massively parallel sequencing and access to the UPPMAX computational infrastructure. We thank M. Li (Sloan Kettering Institute) for *Tgfbr2^fl/fl^* mice.

## Author contributions

H.L. and K.Z. conceived the study with inputs from R.A.H. and H.S.; H.L., K.Z. and R.A.H. designed the experiments. K.Z. performed most of the experiments and analyzed the data under the guidance from H.L. and R.A.H., together with help from H.L. (mice breeding, cell sorting, sample preparation for RNA-sequencing), J.M. (mice breeding and scoring, histopathology), V.J. (single cell library preparation), Y.L. (single cell data analyses), M.P. (mice breeding and scoring), V.S. (cryosectioning, RT-PCR validation), H.S. (behavioral tests, Western blotting). All authors contributed substantially to the final manuscript and discussed the results.

---

**Supplementary Fig 1.**
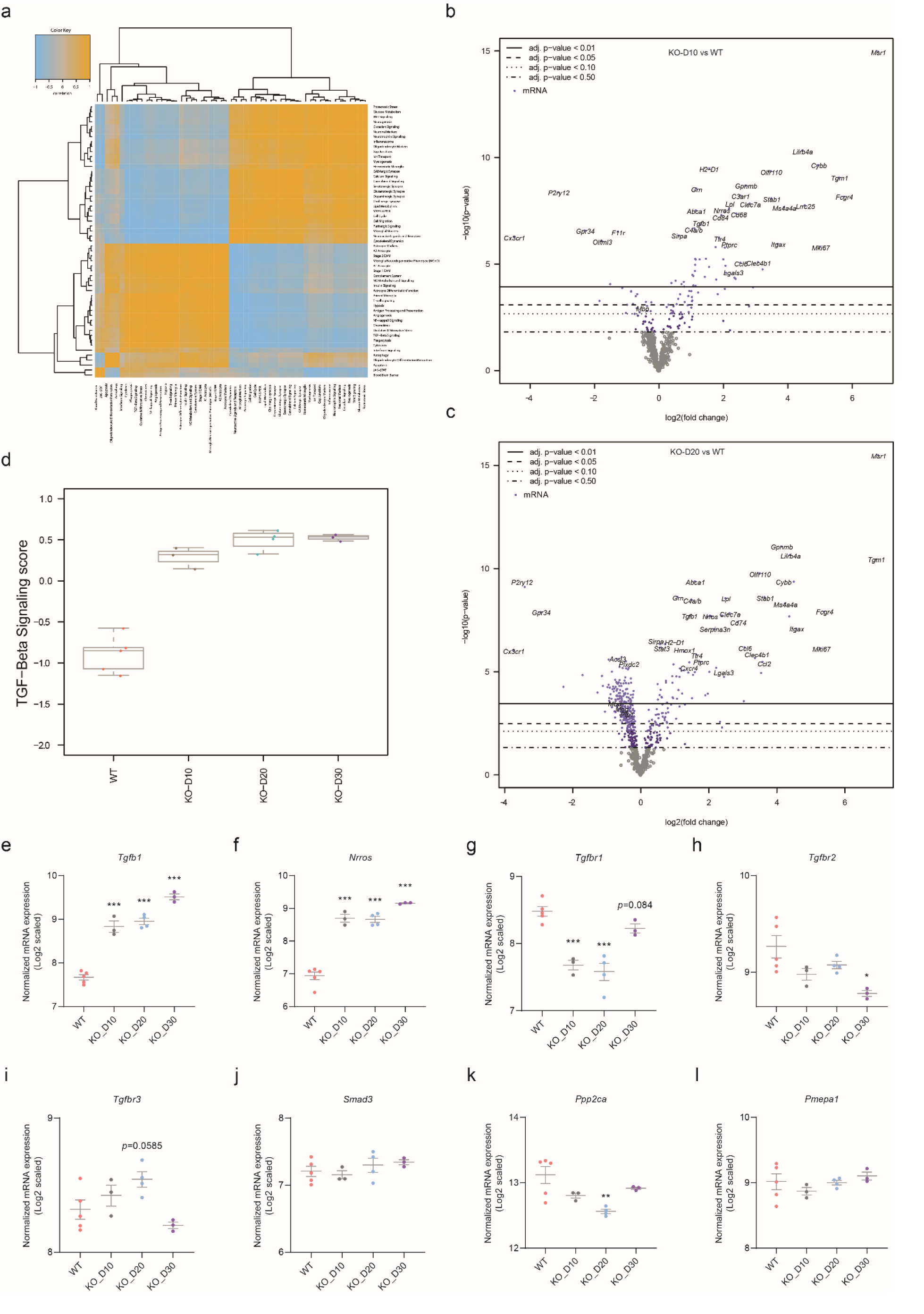
NanoString Glial Profiling of the spinal cord homogenates. **a,** Heatmap showing the correlation matrix of pathway scores. Orange and blue indicate positive and negative correlation, respectively. **b, c** Volcano plots of mRNA count data displaying each gene’s -log10(p-value) and log2 fold change (Tgfbr2^-/-^ KO-D10/WT in panel b, and *Tgfbr2*^-/-^ KO-D20/WT in panel c). Horizontal lines indicate various False Discovery Rate (FDR) thresholds or p-value thresholds if there is no adjustment to the p-values. Genes are colored if the resulting p-value is below the given FDR or p-value threshold. **d,** Chronological pathway score of the TGF-β signaling revealed by NanoString analysis. **e-i,** mRNA expression of genes involved in the TGF-β signaling pathway following microglial *Tgfbr2* deletion. n = 3- 5 biological samples per group. \**p*⍰<⍰0.05; \*\**p*⍰<⍰0.01; \*\*\**p*⍰<⍰0.001 using One-way ANOVA with Dunnett’s Multiple Comparison Test.

**Supplementary Fig 2.**
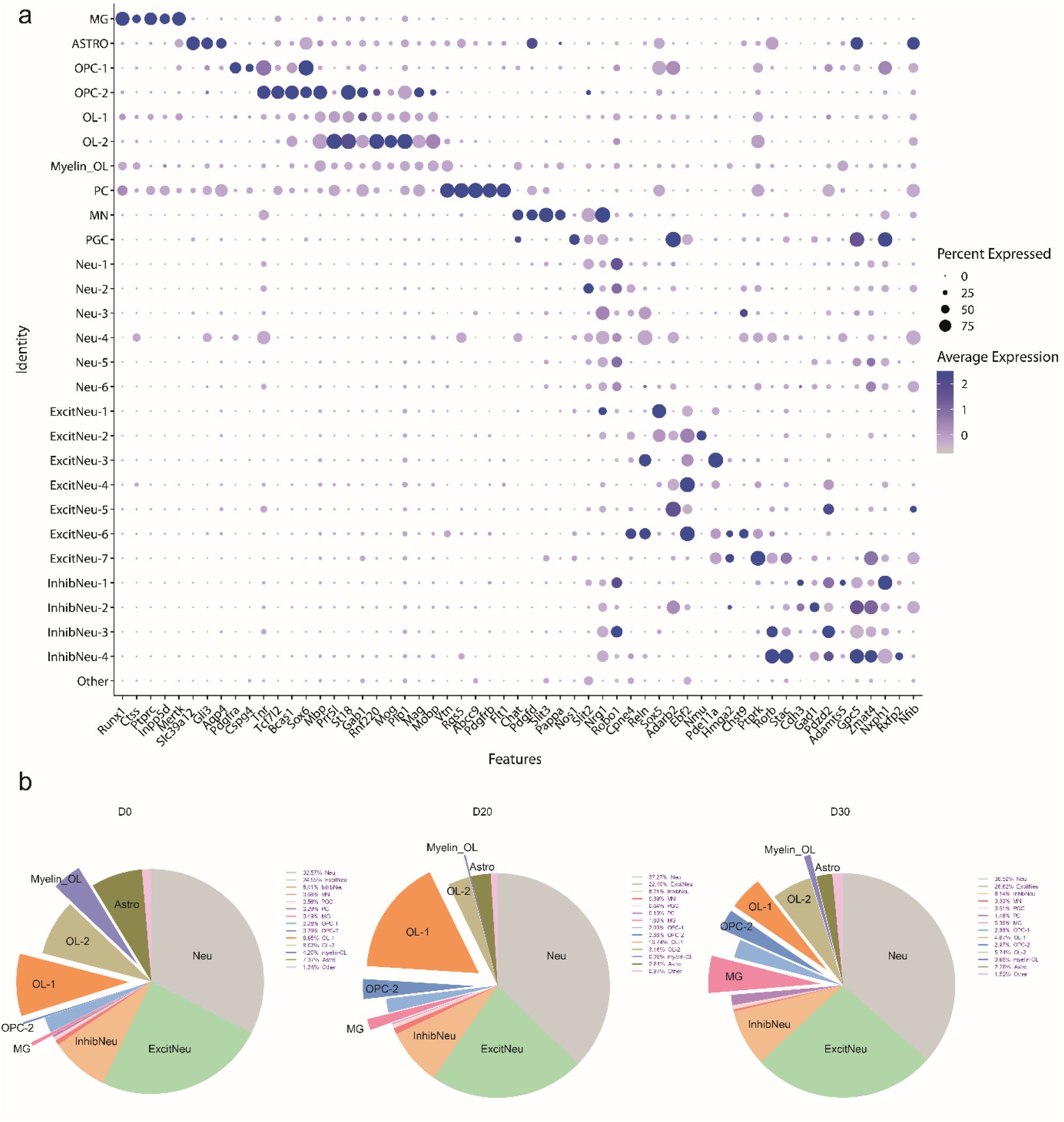
Annotation and cell proportion of the cell clusters revealed in the snRNA-seq. **a,** Bubble plot showing the enriched genes of each cell cluster. **b,** The proportion of the major cell types at different time points.

